# Daily rhythms in the transcriptomes of the human parasite *Schistosoma mansoni*

**DOI:** 10.1101/2021.04.21.440693

**Authors:** Kate A. Rawlinson, Adam J. Reid, Zhigang Lu, Patrick Driguez, Anna Wawer, Avril Coghlan, Geetha Sankaranarayanan, Sarah Kay Buddenborg, Carmen Diaz Soria, Catherine McCarthy, Nancy Holroyd, Mandy Sanders, Karl Hoffmann, David Wilcockson, Gabriel Rinaldi, Matt Berriman

**Author notes:** Corresponding authors: Correspondence to Kate Rawlinson or Matt Berriman.

## Abstract

**Background:** The consequences of the earth’s daily rotation have led to 24-hour biological rhythms in most organisms. Even parasites have daily rhythms, which, when in synchrony with host rhythms, optimize their fitness. Understanding these rhythms may enable development of novel control strategies that take advantage of rhythmic vulnerabilities. Recent work on blood-dwelling protozoan parasites has revealed daily rhythms in gene expression, physiology, drug sensitivity and the presence of an intrinsic circadian clock. However, similar studies on metazoan parasites are lacking. The aims of this study were to investigate if a metazoan parasite has daily molecular oscillations, whether they give us insight into how these longer-lived organisms can survive host daily cycles over a life-span of many years and to determine whether canonical metazoan circadian clock genes are present and rhythmic. We addressed these questions using the human blood fluke *Schistosoma mansoni,* that lives in the vasculature for decades and causes the serious neglected tropical disease schistosomiasis.

**Results:** Using round-the-clock transcriptomics of male and female adult worms we discovered that ∼2% of its genes followed a daily pattern of expression. Rhythmic processes included a night-time stress response and a day-time metabolic ‘rush hour’. Transcriptional profiles in the female reproductive system were mirrored by daily patterns in egg laying (eggs are the main drivers of the host pathology). Genes cycling with the highest amplitudes include drug targets and a vaccine candidate. These 24hr rhythms may be driven by host rhythms and/or generated by a circadian clock. However, core clock genes are missing and orthologues of secondary clock genes show no 24hr rhythmicity in transcript abundance.

**Conclusions:** The daily rhythms identified here reveal temporally compartmentalised internal processes and host interactions over the daily cycle, including processes relevant to within-host survival and between-host transmission. Our findings suggest that if these daily rhythms are generated by an intrinsic circadian clock then the oscillatory mechanism must be distinct from that in other Metazoa. Most importantly, knowing which gene transcripts oscillate at this temporal scale is relevant to functional genomic studies that will lead to the development and delivery of therapeutics against schistosomiasis.

## Background

Most organisms have biological rhythms that coordinate activities with the consequences of the earth’s daily rotation (Reese *et al*., 2017). These biological rhythms are driven by daily cycles in environmental factors such as temperature, light, predation risk and resource availability, as well as an endogenous molecular circadian clock (Rund *et al*., 2016). Whereas some daily phenotypes are driven by natural environmental cycles and become non-rhythmic in constant conditions, circadian rhythms persist in constant conditions, sustained by an endogenous oscillatory mechanism, the circadian clock. The circadian clock is a molecular network that in animals is largely conserved across diverse lineages (Hardin, 2011; Takahashi, 2017). Interconnected regulatory loops are organized around a core transcriptional-translational feedback loop consisting of the positive factors including *CLOCK*, *ARNTL* (*BMAL1/CYCLE*) and the negative regulators *TIMELESS*, *CRYPTOCHROME* and *PERIOD*, and secondary clock genes that modulate the effects of the core feedback loop; together these drive oscillations in many clock-controlled genes. Circadian and clock-controlled genes are a subset of the genes that show daily patterns of expression (diel genes), and all together they lead to 24-hour patterns in physiology and behaviour.

Much like their free-living counterparts, parasites living within the bodies of other organisms also have biological rhythms. These rhythms maximise parasite fitness in terms of within-host survival and between-host transmission (O’Donnell *et al*., 2011). Understanding the rhythms of parasites will provide insight into how they temporally compartmentalise their internal processes and host interactions to survive the daily cycles of host immune system activity and physiology. Furthermore, understanding both parasite and host rhythms may enable development of vaccines and drugs that take advantage of rhythmic vulnerabilities in parasites or harness host rhythms to improve efficacy and reduce drug toxicity (Westwood *et al*., 2019). Recent work on blood-dwelling protozoan parasites has revealed daily rhythms in gene expression, physiology, drug sensitivity (Rijo-Ferreira *et al*., 2017) and the presence of an intrinsic clock (Rijo-Ferreira *et al*., 2020). However, similar studies on metazoan parasites are lacking. Exploring daily molecular oscillations in metazoan parasites will give us insight into how these longer-lived organisms can survive host daily cycles over a life-span of many years, and will lead to an understanding of how animal circadian clockwork has evolved in parasites.

One long-lived metazoan parasite is *Schistosoma mansoni*, a blood-dwelling flatworm (Platyhelminthes), that can live in the vascular system for over 30 years (Harris *et al*., 1984). It causes schistosomiasis, a major neglected tropical disease, that has a profound human impact, with an estimated 140,000 cases and 11,500 deaths in 2019 (GBD 2019). Nothing is known about whether the adult worms (which give rise to the pathology-causing eggs) exhibit any daily or circadian rhythms in any aspect of their biology because they live deep inside the portal veins. Earlier in development *S. mansoni* cercariae larvae are shed from the snail host at a population-specific time of day (Mouahid *et al*., 2012), but the molecular underpinnings of this rhythm are not known. *S. mansoni* naturally infects both humans and mice in the wild (Catalano *et al*., 2018), and mice are commonly used as definitive hosts in the laboratory maintenance of its life cycle. The mouse is also the model species routinely used to study circadian rhythms in mammals (Zhang *et al*., 2014), and because of this, we know there are many daily rhythmic fluctuations in the vasculature (e.g. temperature, pressure, oxygen, glucose, red and white blood cells [Damiola *et al*., 2004, Curtis *et al*., 2007, Scheiermann *et al*., 2012, Adamovich *et al*., 2017, Llanos & de Vacarro, 1972]) that may act as zeitgebers (German for “time giver” or synchronizer) to influence the worm’s biology and potentially its rhythms. Here we ask whether sexually mature male and female *S. mansoni*, collected from their natural environment (the mesenteric vasculature of mice) under ‘normal’ host conditions (Light:Dark cycle) show daily rhythms in their transcriptomes. Transcripts that cycle with 24 hour periodicity (diel or cycling genes), were identified using RNA-seq time series from female and male worms as well as the heads of males (to enrich for the worm’s cephalic ganglia or ‘brain’; the site of the master circadian clock in some animals). Protein function databases and published single cell RNA-seq data (Wendt *et al*., 2020) were interrogated using our diel genes to identify rhythmic biological processes. One-to-one orthologs were compared between *S mansoni* diel genes and those of other animals to investigate metazoan circadian clock components. Having found that most cell types associated with the female reproductive system are enriched for cycling genes, we show that egg laying *in vitro* oscillates between day and night. Our discovery that genes oscillate throughout the 24-hour period has given us an understanding of the fine-scale temporal partitioning of biological processes in male and female worms, and indications of parasite/host interactive rhythms. As these diel genes include potential drug targets and a vaccine candidate, this study will benefit the development and delivery of treatments against schistosomiasis.

## Results

We collected mature male and female worms at four hour intervals over a 44 hour period from mice entrained in alternating 12-hour light and dark cycles (LD12:12). Although light is probably not a relevant zeitgeber to schistosome adults (as penetrative light levels below 5mm in mammalian bodies are minimal, Ash *et al*., 2017), light is an important cue for the mouse, and as nocturnal creatures, they are active mainly during the dark phase (Jud *et al*. 2005). Zeitgeber time (ZT) 0 indicates the beginning of the light phase and resting phase for the mouse (which we call day-time here), and ZT 12 is the beginning of dark phase and active phase for the mouse (called night-time in this study). Male and female worms from each mouse were separated and pooled. Heads of male worms were also isolated and these were pooled for each mouse. RNA was extracted from each pool and sequenced. This gave us three time-series datasets; one for whole female worms, one for whole males and one for male heads (Fig. 1A).

**Fig. 1.**
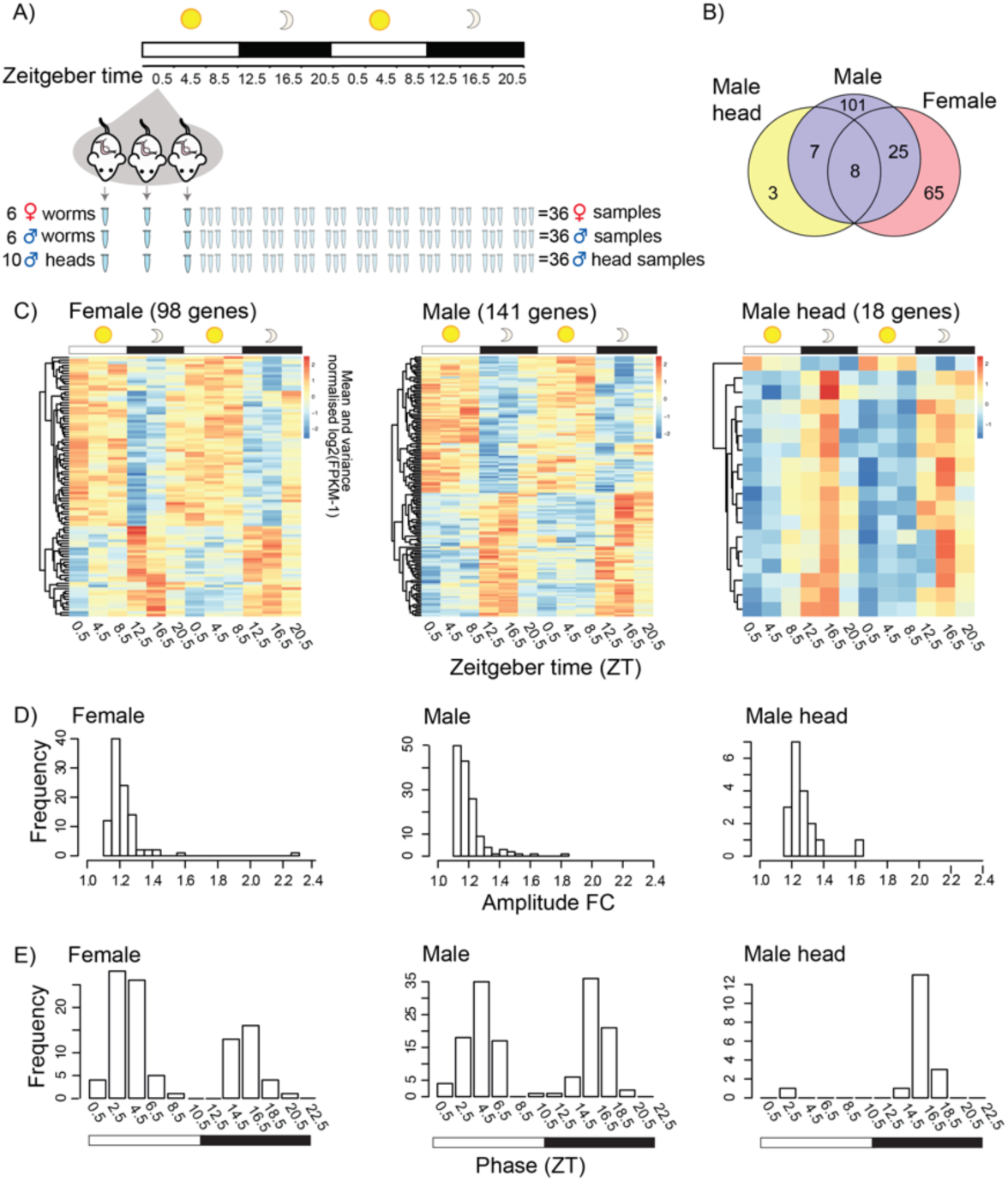
*Schistosoma mansoni* genes with 24-hour periodicity in their expression in adult worms. **A**) Schematic of the collection of pooled worm samples every 4 hours over 44 hours. **B**) Overlap in 209 diel genes between females, males and male heads. **C**) Expression heatmaps of diel genes. Each row represents a gene whose transcripts oscillate with ∼24 hour periodicity, ordered vertically by phase. **D**) Histograms of rhythmic daily fold changes (FC) in transcript abundance. **E**) Histograms showing bimodal peaks of expression in diel genes in both sexes, but in male heads most diel genes peak at midnight.

### 1. Daily rhythms in the transcriptomes of adult *Schistosoma mansoni*

From each dataset, we identified genes that were differentially expressed (FDR < 0.05) over a 24 hour period: 206 in females, 194 in males and 48 in male heads (Additional file 1: Table S1-3). We then determined which differentially expressed genes were oscillating with a periodicity close to 24 hours using JTK_cycle (Hughes et al., 2010), and found 98 diel genes in females, 141 in males and 18 in male heads (Fig. 1B&C; Additional file 1: Table S1-4). A significant number were shared between males and females (33; Fisher’s exact test odds ratio = 47, *P* < 10^-16^) and between males and male heads (15; Fisher’s exact test odds ratio = 395, *P* < 10^-16^; Fig. 1B, Additional file 1: Table S4). The combined number of diel genes from the three datasets was 209 (Fig. 1B; Additional file 1: Table S4), 1.9% of the *S. mansoni* (v7) gene set. The median peak-to-trough fold change (amplitude) of gene expression for diel genes was 1.19, 1.18 and 1.24 for females, males and male heads respectively. However, many genes had much higher daily fold-changes in expression (Fig. 1D; Table 1); the highest in each of the datasets were *SmKI-1* (Smp_307450, fold change 1.8) in males that encodes a BPTI/Kunitz protease inhibitor domain protein, as well as *hsp70* (Smp_049550, 1.6 fold) in male heads and *hsp90* (Smp_072330, 2.3 fold) in females that each encode heat shock proteins (in males *Hsp90* has a 4.8 fold change but falls just outside FDR <0.01 threshold [FDR = 0.0103])(Table 1)(see Additional file 1: Table S5 for orthologs in other taxa). While we identified rhythmic genes with peaks of expression at most times of day, there was a clear bimodal pattern in both females and males (Fig. 1E). In females, expression peaked at 2.5- 4.5hr (ZT) after lights on and at 14.5-16.5 hr (ZT) (2.5-4.5hr after lights off), but with 66% of the cycling genes peaking during the day (the hosts resting phase), while in males the peaks occurred at 4.5 and 16.5 (ZT) (with a 53:47% split between light:dark phases). In the heads of males, the expression of diel genes peaked at ZT 16.5, and was not bimodal, with all but one reaching peak expression during the night (the hosts active phase).

**Table 1.** Diel genes with the highest expression amplitudes. Daily fold-changes in transcript abundance are shown for female and male worms, plus male heads, with peak phase of expression (Zeitgeber Time, ZT, bold = night) and significance of 24-hr rhythmicity determined using JTK package. (* I-Tasser predicted structural analogue, see Additional file 1: Table S6).

| Dataset | Gene ID | Gene description | Fold change | Phase (ZT) | JTK Benjamini-Hochberg q-value |
| --- | --- | --- | --- | --- | --- |
| Female | Smp_072330 | Heat shock protein 90 | 2.30 | **14.5** | 0.00032 |
|  | Smp_326610 | Trematode eggshell synthesis domain-containing protein | 1.56 | 6.5 | 0.00191 |
|  | Smp_004780 | FKBP-type peptidylprolyl isomerase (PPIase) | 1.41 | **16.5** | 0.00019 |
|  | Smp_044850 | Ribokinase | 1.41 | **16.5** | 0.00770 |
|  | Smp_342000 | FMRFamide-activated amiloride-sensitive sodium channel | 1.38 | 8.5 | 0.00191 |
|  | Smp_069130 | Heat shock protein 70 (Hsp70)-4 | 1.35 | **14.5** | 0.00029 |
|  | Smp_319380 | n/a | 1.33 | 2.5 | 0.00362 |
|  | Smp_244190 | ZP domain-containing protein | 1.31 | **14.5** | 0.00588 |
|  | Smp_340010 | Putative eggshell protein | 1.30 | 4.5 | 0.00235 |
|  | Smp_064860 | Putative heat shock protein 70 (Hsp70)-interacting protein | 1.30 | **16.5** | 7.92E-05 |
| Male | Smp_307450 | SmKI-1, a BPTI/Kunitz protease inhibitor domain protein | 1.83 | 4.5 | 0.00311 |
|  | Smp_049550 | Putative heat shock protein 70 (Hsp70) | 1.62 | **16.5** | 1.21E-06 |
|  | Smp_324960 | Hypothetical protein (Fatty acid synthase-like*) | 1.57 | **18.5** | 0.00112 |
|  | Smp_327270 | Very-long-chain 3-oxoacyl-CoA reductase | 1.52 | **18.5** | 0.00763 |
|  | Smp_327260 | Transmembrane protein 45B | 1.49 | **18.5** | 0.00468 |
|  | Smp_013950 | Solute carrier family 43 member 3 | 1.48 | 0.5 | 0.00112 |
|  | Smp_328570 | n/a | 1.48 | 4.5 | 0.00763 |
|  | Smp_327240 | Hypothetical protein (GTPase activator activity*) | 1.45 | **18.5** | 0.00651 |
|  | Smp_004780 | FKBP-type Peptidylprolyl isomerase (PPIase) | 1.36 | **16.5** | 7.31E-05 |
|  | Smp_013790 | Probable ATP-dependent RNA helicase, DDX5 | 1.35 | 0.5 | 0.00047 |
| Male head | Smp_049550 | Putative heat shock protein 70 (Hsp70) | 1.63 | **16.5** | 1.43E-05 |
|  | Smp_004780 | FKBP-type peptidylprolyl isomerase (PPIase) | 1.35 | **16.5** | 0.00055 |
|  | Smp_145560 | Monocarboxylate transporter 9 | 1.33 | 2.5 | 0.00236 |
|  | Smp_124820 | Putative chromosome region maintenance protein 1/exportin | 1.30 | **18.5** | 0.00236 |
|  | Smp_069130 | Heat shock protein 70 (Hsp70)-4 | 1.30 | **16.5** | 0.00236 |
|  | Smp_333170 | 85/88 kDa calcium-independent phospholipase A2 | 1.28 | **18.5** | 0.00188 |
|  | Smp_113620 | Serine/arginine-rich splicing factor 2 | 1.27 | **16.5** | 0.00297 |
|  | Smp_049600 | Putative DNAj (Hsp40) homolog, subfamily C, member 3 | 1.25 | **16.5** | 7.31E-05 |
|  | Smp_214080 | VEZF1/Protein suppressor of hairy wing/Zld/ C2H2-type | 1.25 | **16.5** | 7.49E-06 |
|  | Smp_246230 | Nicotinate-nucleotide pyrophosphorylase | 1.24 | **16.5** | 0.00035 |

### 2. Putative functions of diel genes

To better understand the possible function of the diel genes, we used a combination of annotation-enrichment analyses based on Gene Ontology (GO), KEGG pathways, STRING molecular interactions and the adult *S. mansoni* single-cell RNA-seq dataset from Wendt *et al*. (2020). Our analyses show that the diel genes are involved in distinct rhythmic processes during the night and day time.

#### Night-time peaking genes

Of the 63 genes with a night-time peak in males, 33 formed a single large network based on their predicted STRING interactions (Fig. 2). Within this network many of the molecules with the largest number of interactions were putative molecular chaperones (orthologs to heat shock proteins [HSPs] and their co-chaperones) involved in protein processing the ER and the stress response. In females a smaller network of 13 genes was predicted, but heat shock /stress response genes were still the main constituents (Additional file 2: Fig. S1). These observations were supported by enrichment of GO terms related to protein folding and chaperones: protein folding/unfolded protein binding (males FDR 0.0013, females FDR 10^-9^; Additional file 1: Table S7). Diel genes involved in these processes and networks all had acrophases between 14.5-18.5 ZT (i.e. mid dark phase)( Additional file 3: Fig. S2), and could be mapped to three KEGG pathways; ‘protein processing in the endoplasmic reticulum’ (Additional file 4: Fig. S3), ‘PI3K-AKT signalling’ (Additional file 5: Fig. S4) and ‘Estrogen signalling’(Additional file 6: Fig. S5). Orthologs of six of the diel HSPs in *S. mansoni* show 24-hour rhythms in other animals as well (Additional file 1: Table S4, 5 & 13).

**Fig. 2.**
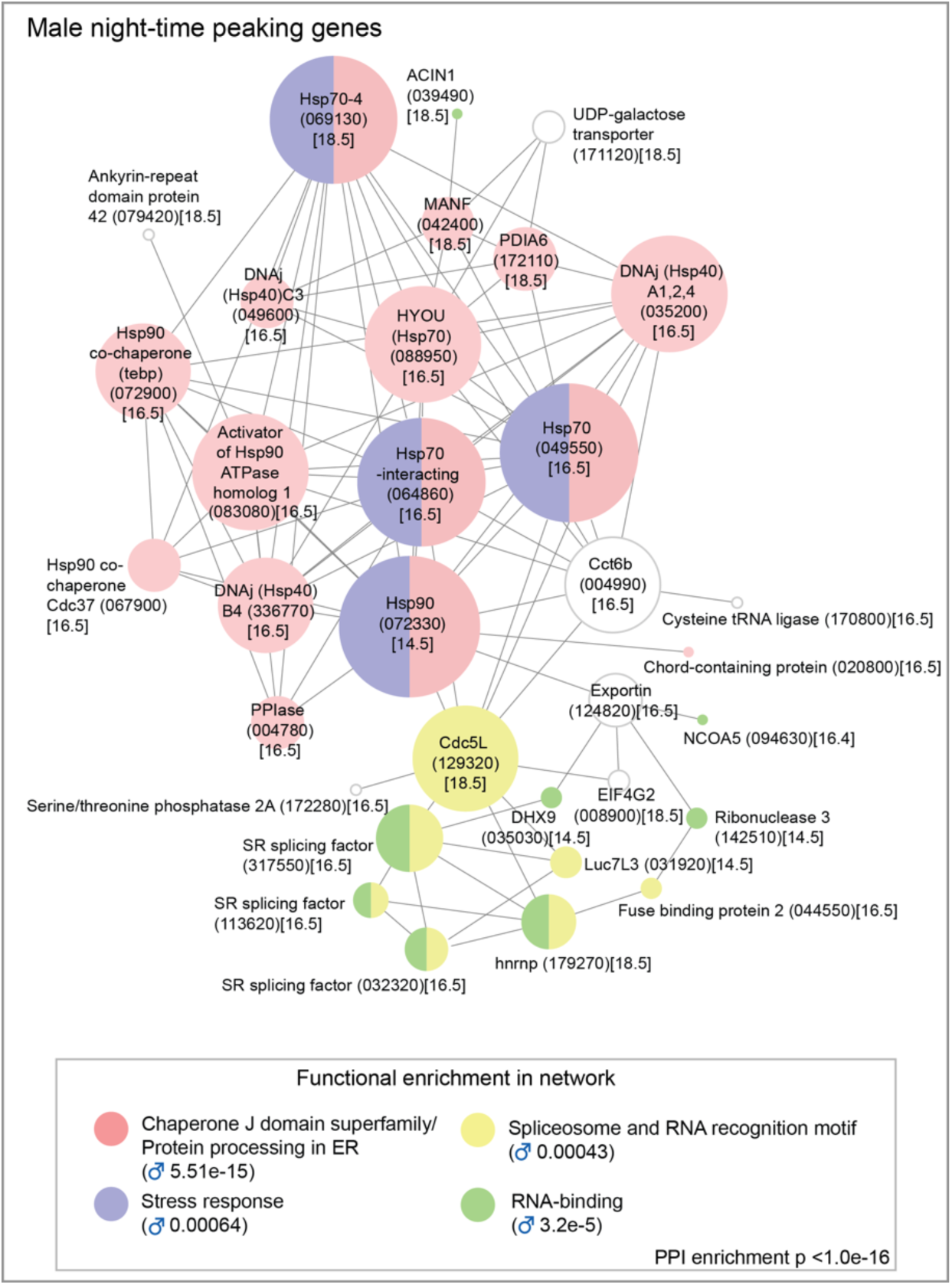
Predicted molecular interaction networks of night-time peaking genes in male *Schistosoma mansoni* (computed using the STRING online database). All genes peaked between 14.5-18.5 ZT (i.e. between 2.5- 6.5 hours after the start of the dark phase). Node size reflects the number of connections a molecule has within the network. Lines (edges) connecting nodes are based on evidence of the function of homologues. Functional enrichment (FDR) as provided by STRING. (PPI= predicted protein interaction; gene identifiers shown in parenthesis but with “Smp_” prefix removed for clarity; timing of acrophase (ZT) in square brackets).

Ten HSPs and co-chaperones cycled in both sexes, with eight oscillating in phase synchrony between the sexes (Additional file 3: Fig. S2). A striking example of this is seen in *hsp90* (Smp_072330) and *FKBP-type peptidylprolyl isomerase (PPIase)*(Smp_004780); *hsp90* peaks at the start of the dark phase, 4 hours before *PPIase* (Fig. 3 & 4). In other animals, these proteins form part of a heterocomplex that chaperones steroid hormones in cell signalling (Additional file 6: Fig. S5) and they are critical for reproductive success (Cheung-Flynn et al., 2005). Although their functions aren’t known in *S.mansoni*, both are single cell markers for germ stem cell progeny and we show, using whole mount *in situ* hybridisation (WISH), that *PPIase* is expressed in sperm throughout the testes and in all oocytes in the ovary (Fig. 3C & 4C). *PPIase* also cycles in male head samples and is expressed in many cells in the head (Fig. 3C). There were also sex differences in the diel genes involved in the nightly chaperone response. For example, a *DNAj (hsp40*), which has homologs located on the different sex chromosomes (known as gametologues)(Buddenborg *et al*., *in prep*), has diel expression in both gametologues. Schistosomes have a ZW sex chromosome system, where females are ZW and males are ZZ. The Z gametologue (Smp_336770) cycles in females, males and male heads, and the W gametologue (Smp_020920) also cycles, and is present only in females. Further sex differences included diel expression, in males only, of nine genes whose orthologs are involved in stress response and recovery (Additional files 3: Fig. S2).

**Fig. 3.**
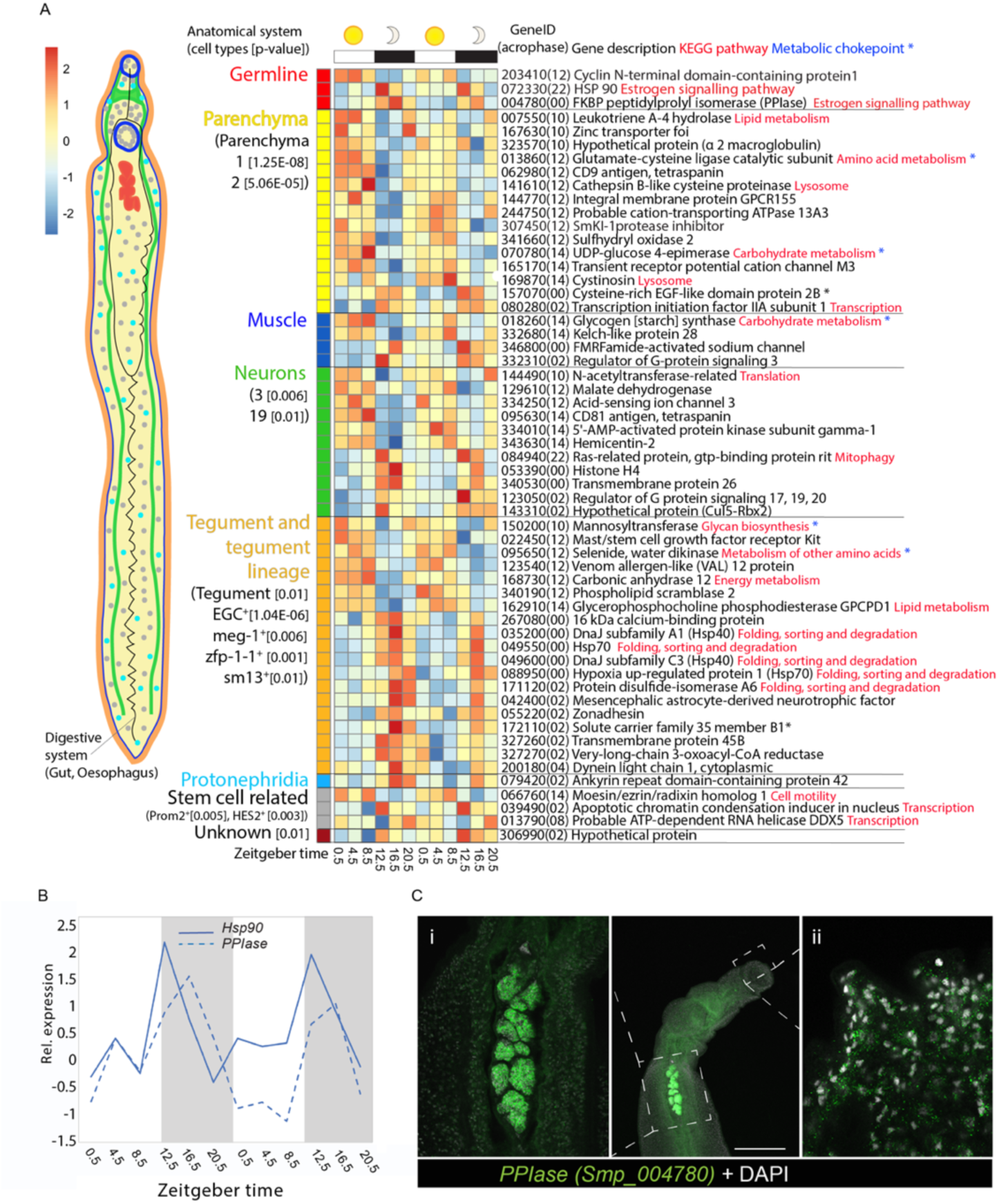
A) Heatmap of the 57 male diel genes that are single cell markers (identified in Wendt *et al*., 2020). Each row represents a diel gene, ordered vertically by phase within anatomical systems containing constituent cell types. Diel genes that are markers for multiple cell type categories are placed within cell type (and their category) with greatest difference between the Seurat pct1-pct2 scores. “Smp_” prefixes have been removed from gene identifiers for clarity; timing of acrophase in parentheses. P-value of cell types significantly enriched in diel genes given in brackets [p<0.05]. **B)** Temporal relative expression profiles of *Hsp90* (Smp_072330) and *FKBP-type peptidylprolyl isomerase (PPIase)*(Smp_004780) showing peaks of expression at night. **C**) WISH expression of *PPIase* showing labelled transcripts in male worm i) throughout the testes, and ii) in the head (scale bar = 100µm)(100% of individuals examined, n = 10).

Four of these genes are markers for tegument-related cell types (Fig. 3) and as the male tegument forms a large surface area that is in direct contact with mouse blood and endothelium, these cells are likely to be exposed to environmental heat shock triggers. Night-time peaking genes were also associated with GO terms related to RNA binding and mRNA splicing, but in males only (FDR = 0.0002 for RNA binding, FDR=0.0292 for mRNA splicing; Additional file 1: Table S7). Based on STRING, many of these genes form part of the night-time network (Fig. 2), including putative homologues of: *human Ser/Arg-rich splicing factors* (Smp_317550, Smp_032320, Smp_113620); *heterogeneous nuclear ribonucleoprotein* (Hnrp; Smp_179270): the *cell division cycle control protein Cdc5L* (Smp_129320), a spliceosome component; and the splicing regulator (Li *et al*., 2013) *far upstream element-binding protein* (Smp_044550). The night-time network predicts an interaction between heat shock proteins and RNA-binding and mRNA splicing genes, connected via *Cdc5L* (Fig. 2).

Diel genes involved in regulation of GPCR functioning (*regulator of G-protein signalling 3 [RGS3]* Smp_332310; and *RGS20* Smp_123050) peaked at night and are markers for muscle and nerve cells respectively (Fig. 3 and 4). *Histone 4* (Smp_053390) and *gtp-binding Ras-related protein* (Smp_084940)(a member of the histone co-chaperone pathway) also peaked at night in males and are markers for nerve cell types (Fig. 3). Neuronal activity promotes histone turnover (Maze *et al*., 2015; Grover *et al*., 2018) and taken together these results may indicate higher sensory responsiveness and activity night.

**Fig. 4.**
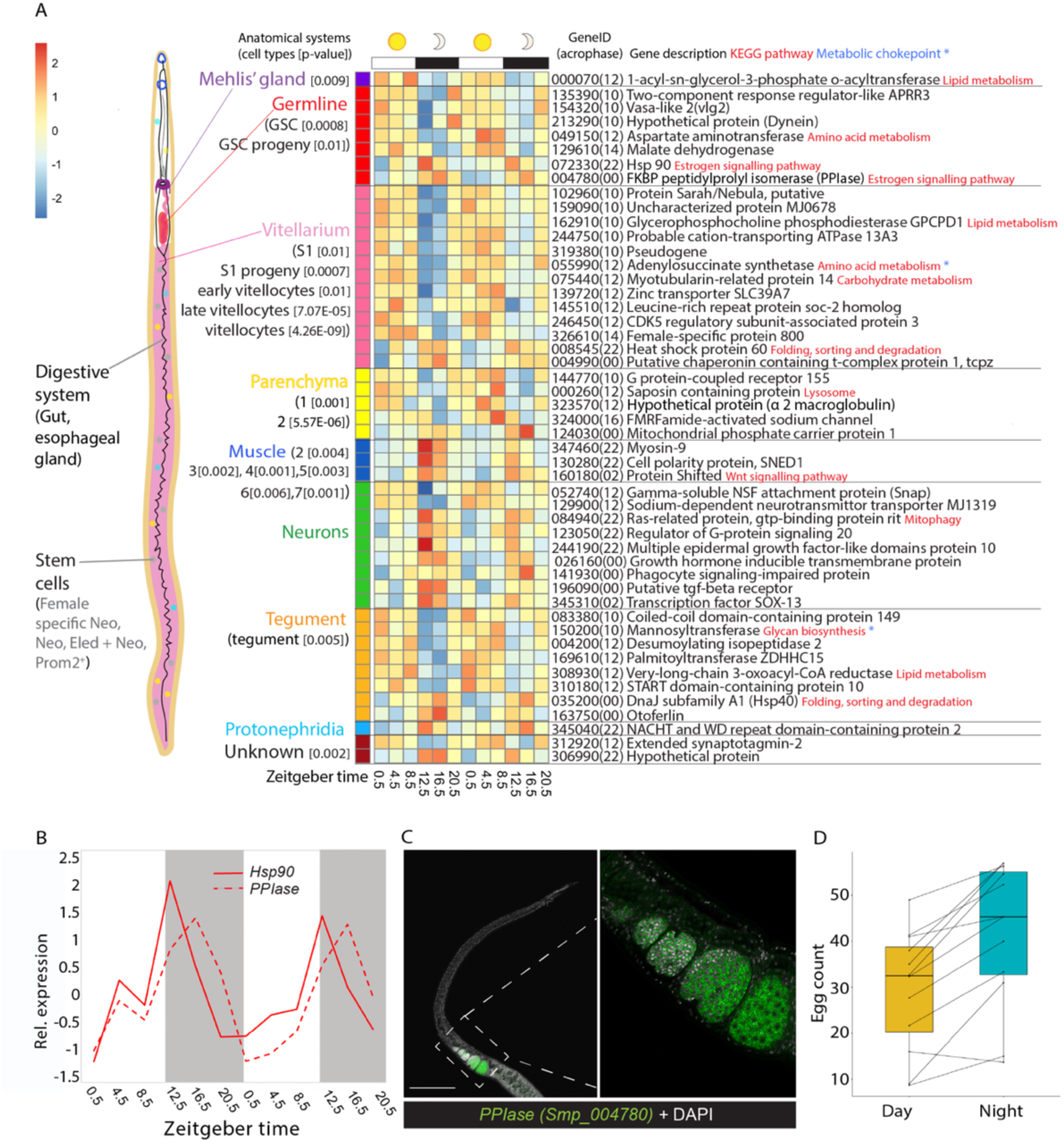
A) Heatmap of 48 single-cell marker genes that showed diel expression in females. Marker genes are from Wendt et al., 2020. Each row represents a diel gene, ordered vertically by phase within anatomical systems containing constituent cell types. Diel genes that are markers for multiple cell type categories are placed within cell type (and their category) with greatest difference between their Seurat pct1-pct2 scores. “Smp_” prefixes have been removed from gene identifiers for clarity; timing of acrophase in parentheses. P-value of cell types significantly enriched in diel genes given in brackets [p<0.05]. B) Temporal relative expression profiles of *Hsp90* (Smp_072330) and *FKBP-type peptidylprolyl isomerase (PPIase)*(Smp_004780) showing peaks of expression at night. C) WISH expression of *PPIase* showing labelled transcripts in female worm in all oocytes in the ovary (scale bar = 100µm)(100% of individuals examined, n = 10). D) Female (paired) worms *in vitro* lay more eggs at night (median and interquartile ranges) than during the day (n=12 female worms; median (night egg count - day egg count) = 12.0; paired Wilcoxon test: P=0.003216).

#### Day-time peaking genes

Predicted interactions between the day-time peaking genes were more limited than at night; both sexes had interactions of genes associated with metabolism (Additional file 7: Fig. S6). There were 32 diel genes that could be mapped to KEGG metabolic pathways; most of which are involved in lipid, carbohydrate and amino acid metabolism (Additional file 1: Table S9). Of these, 28 have acrophases between 0.5–6.5 ZT, suggesting a peak in metabolic activity at this time; an extended metabolic ‘rush hour’ (Fig. 5A). We identified twenty-one diel genes that have previously been classified as metabolic chokepoints (capable of uniquely generating specific products or utilising specific substrates, International Helminths Genome Consortium, 2019) for potentially targeting with new drugs, 15 of these peaked during the day (Fig. 5B) and six at night (Additional file 1: Table S4). Several metabolic diel genes were also markers of specific cell types: those of the reproductive system in female worms, and those of tegumental, parenchymal and muscle cells in males (Fig. 3 & 4).

**Fig. 5.**
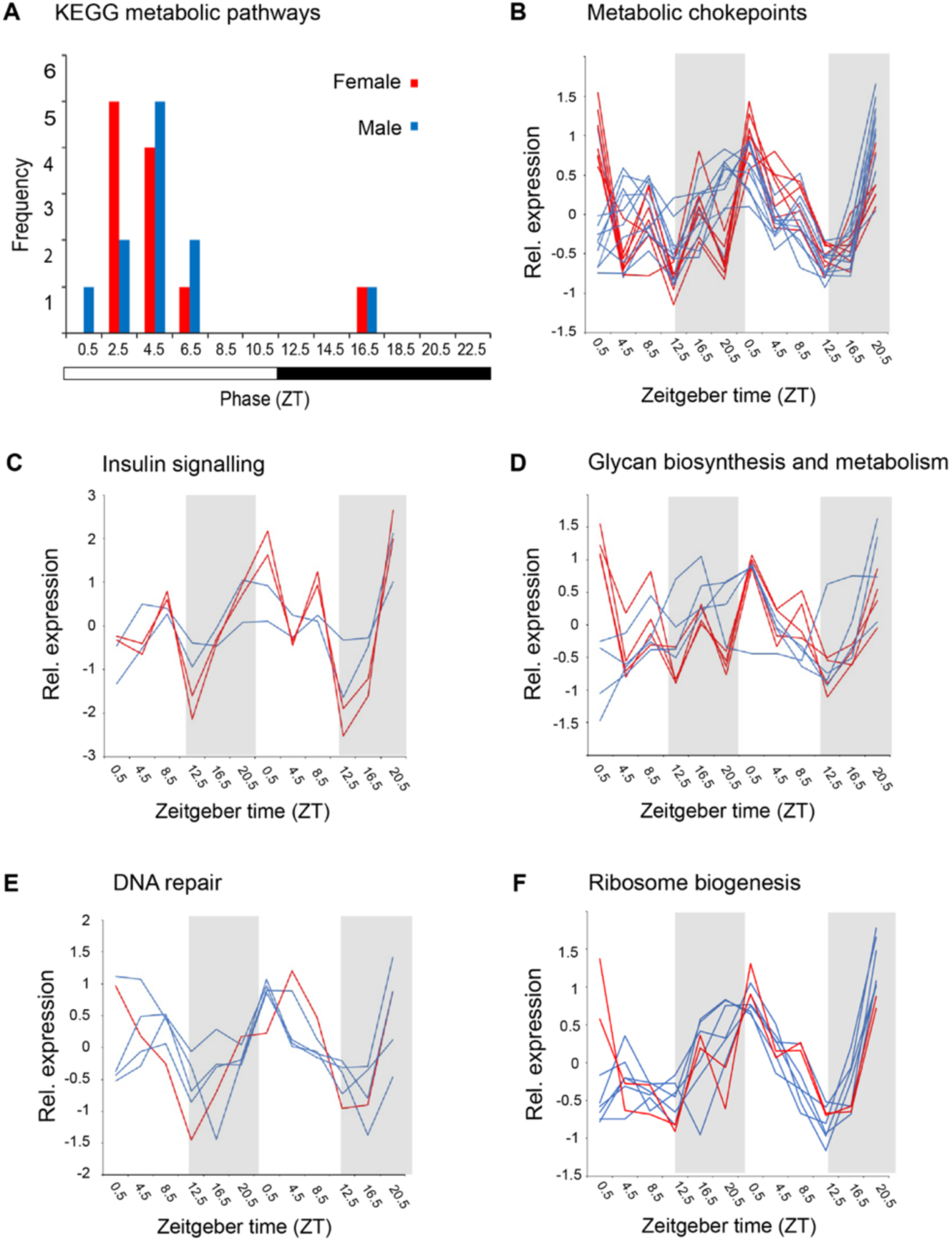
Light-phase peaking diel genes are involved in metabolism, DNA repair and ribosome biogenesis. **A**) Peak phase of expression of diel genes in KEGG pathways for metabolism. Temporal expression profiles of **B**) day-time peaking metabolic genes identified as chokepoints and potential drug targets, **C**) genes involved in the insulin signalling pathway **D**) genes involved in glycan biosynthesis and metabolism, **E**) DNA repair and **F**) ribosome biogenesis.

There are four diel genes involved in the insulin signalling pathway and they all peaked between 2.5-6.5 ZT; *glycogen synthase* (Smp_018260) and *3-phosphoinositide-dependent protein kinase 1* (Smp_094250) in males, and *hexokinase* (Smp_043030) and *protein phosphatase 1 regulatory subunit 3B* (Smp_167660) in females (although the latter three fell between FDR adjusted p-values of 0.01-0.05)(Additional file 1: Table S9, Fig. 5D).

Glycans and glycoproteins that are secreted or localized to the tegument interact with the host immune and hemostatic systems (Smit *et al*., 2015). We found six diel genes involved in N-glycan and glycosaminoglycan synthesis, and glycosylation (Mickum *et al*. 2014), five of which peaked during the day (Fig. 5D, Additional file 1: Table S9), including mannosyltransferase (Smp_150200) that is a marker for a tegumental cell type (Fig. 3 & 4; Additional file 1: Table S8). Three others encode enzymes involved in synthesis of heparin-like glycosaminoglycans that may increase anti- coagulation activity of mammalian host blood (Mebius *et al*., 2013); putative beta-1,3- glucuronyltransferase (heparin-like)(Smp_083130) and Zinc finger CCHC domain-containing protein 4 (Smp_245920) peak during the day in females, whereas putative heparan sulfate n-deacetylase/n-sulfotransferase (Smp_134250) cycles in males, but peaks at night (Additional file 1: Table S9).

Also peaking during the day are other genes involved in host-parasite interactions. *SmKI-1* (Smp_307450) is one of four similar genes that correspond to Smp_147730 from an earlier version (v5) of the genome assembly (see Additional file 8: Supplementary information 1). The SmK1-1 protein is localised to the tegument and secreted into the host where it inhibits host proteases (including neutrophil elastases) involved in triggering the immune response (Morais *et al*. 2018), and interferes with host coagulation pathways to delay blood clot formation (Ranashige *et al*., 2015). Smp_307450 reaches its peak at midday (ZT 4.5)(Fig. 6) a few hours before the daily increase of mouse blood coagulation factors (Bertolucci *et al*., 2005) and coinciding with the day- time release of mouse neutrophils into the blood from bone marrow (De Filippo & Rankin, 2018), possibly indicating that its diel expression may be anticipating or responding to these host cues.

**Fig. 6.**
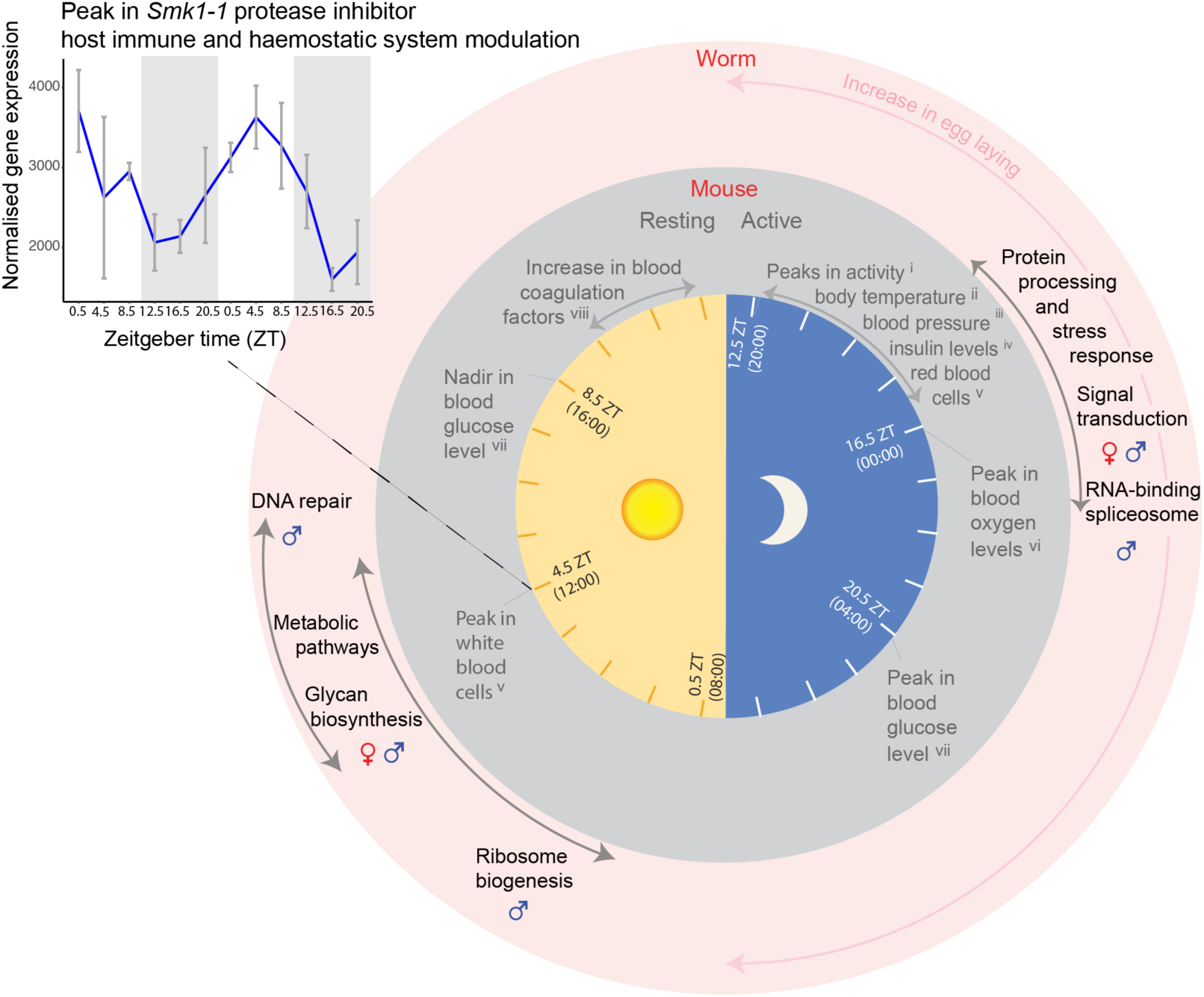
Summary schematic representing the daily rhythms in the transcriptomes of adult *Schistosoma mansoni* (from diel gene enrichment analyses) and the mouse vasculature (^i^Jud *et al*., 2005, ^ii^Damiola *et al*., 2004, ^iii^Curtis *et al*., 2007, ^iv^Feillet *et al*., 2016, ^v^Scheiermann *et al*., 2012, ^vi^Adamovich *et al*., 2017, ^vii^Llanos & de Vacarro, 1972, ^viii^Bertolucci *et al*., 2005). Side plot showing temporal expression profile of *Smk1-1*(Smp_307450) in males. Exponentiated, VST-normalised gene expression values are shown, with means taken for each timepoint. Error bars show the standard error of the mean.

Also peaking at 4.5 ZT are *carbonic anhydrase 12* (Smp_168730), which is a glycoprotein localised on the surface of tegument and contributes to parasite survival and virulence (Da’dara *et al*., 2019), and *Val12* (Smp_123540) which is a putative secreted member of the venom allergen-like family (Chalmers *et al*., 2008). Both are single cell markers for tegumental cell types. *Val12* is a marker for late tegumental progenitor cells (Sm13^+^ cells)(Wendt *et al*., 2018), and we show it expressed in approximately 500 cells that are positioned along the entire length of the male body (Additional file 9: Fig. S7). *Val12^+^* cells are more abundant dorsally than ventrally, and transcripts reach the body wall musculature, with some extending in the tubercles of the dorsal tegument (Additional file 9: Fig. S7). This *in situ* expression supports protein structural data indicating that it is likely to be secreted/excreted, onto the tegument or into the host environment (Chalmers *et al*., 2008).

In males two additional day-time networks of interacting genes were predicted. The first was formed of four genes involved in DNA repair (Additional file 7: Fig. S6), and the GO term ‘damaged DNA binding’ was significantly enriched (FDR = 0.0036, Additional file 1: Table S7; Additional file 10: Fig. S8). The second included genes putatively involved in ribosome biogenesis (Additional file 7: Fig. S6); a putative DEAD box ATP-dependent RNA helicase (orthologous to DDX5)(Smp_013790) which cycles with high amplitude (1.4) and is a marker of HES2+ stem cells (Fig. 3), and a putative AAA family ATPase (Smp_160870) (Fig. 2) which also exhibits diel expression in females, and whose human ortholog (NVL) regulates 60S ribosomal subunit biogenesis in the nucleolus in a spatiotemporal manner (Nagahama *et al*., 2004).

The transcriptome patterns associated with the female reproductive system are particularly interesting because schistosome eggs are the cause of disease in the host (including granuloma formation after becoming embedded in the liver and intestine) and transmission. Seventeen of the 21 diel genes that are markers for cell types of the female reproductive system (the vitellarium, germline and Mehlis’ gland cells) peak during the day, between 2.5-6.5 ZT. These included a Trematode Eggshell Synthesis domain-containing protein (Smp_326610; Table 1, Fig. 4A, Additional file 1: Table S10) that is one of several major structural component of the egg shell in schistosomes (Ebersberger *et al*., 2005). Also expressed in phase are other diel genes potentially involved in reproduction (but not identified as cell type markers); a *vasa-like DEAD box ATP-*

*dependent RNA helicase* (Smp_154320, expressed in mature oocytes, *Smvlg2* [Skinner *et al*., 2012]), and a second *putative eggshell protein* (Smp_340010). To investigate whether transcriptional rhythms might reveal patterns in egg laying, we recorded day and night egg counts for individual worm pairs *in vitro*. For each of 12 worm pairs, we calculated a day egg count as the median of three consecutive days (three replicates), and a night egg count as the median of three consecutive nights (three replicates). Across the 12 worm pairs, the difference between the night egg count and the corresponding day egg count was significantly higher than zero, that is, paired females tended to lay more eggs at night than during the day (n=12 females; median (night egg count - day egg count) = 12.0; paired Wilcoxon test: P=0.003) (Fig. 4D). This was also seen in three further independent biological replicates (replicate 2: n=17, paired Wilcoxon test P=0.002, median(night-day)=5.3, replicate 3: n=17, P=0.0003, median(night-day)=17.5; replicate 4: n=8, P=0.008, median(night-day)=14.3) (Additional file 11: Fig. S9). Females, on average, laid 13 more eggs over the 12-hour dark phase, corresponding to a 50% increase compared with the light phase.

### 3. Could daily rhythms be generated by a circadian clock intrinsic to *Schistosoma mansoni?*

Of the 33 genes that cycle in both sexes, 17 show identical phases and the remaining show acrophases within 4hrs of the opposite sex (Additional file 12: Fig. S10; Additional file 1: Table S4). These mRNA oscillations in phase between the sexes suggest external cyclical cues drive the synchrony of these rhythms within the *S. mansoni* pairs (and population) inside the mammalian host. Host rhythms may be driving the parasite rhythms, and doing so either directly, or entraining an endogenous time-keeping machinery within the parasite – an intrinsic circadian clock.

This led us to investigate if *S. mansoni* has homologues of the canonical animal circadian clock genes and whether they show 24-hour periodicity in our datasets. Extensive BLASTP searches revealed that core elements of the negative feedback loop appear to be missing. Even when the stringency was relaxed to an E-value of 1 (BLAST), there were no putative hits for Period or any Cryptochromes (Additional file 1: Table S11). We found two DNA photolyases (Additional file 1: Table S12), which are members of the cryptochrome/photolyase family (CPF), but they clustered with the CPD photolyases, not the canonical animal circadian-related Cryptochromes (Additional file 13: Fig. S11) and they lacked the FAD binding domain (see Additional file 14: Supplementary information 2) for secondary structural features of putative circadian proteins). We identified three bHLH-PAS proteins similar to Clock (Smp_178780 and Smp_168600) and Cycle/Bmal1 (Smp_341950)(Additional file 1: Table S11 & 12), however phylogenetic analysis showed that they clustered with the closely-related non-circadian proteins ARNT, AHR and SIM (Additional file 13: Fig. S11). Therefore, *S. mansoni* appears to lack the core negative feedback genes, Period and Cryptochromes, as well as the positive transcription factors Clock and Bmal1/Cycle. However, we did identify an orthologue for Timeout (Tim2)(Smp_163340), but not its paralogue Timeless (Tim1) (Additional file 13: Fig. S11, Additional file 1: Table S11 & 12). *Timeout* doesn’t cycle in our datasets.

We identified putative homologs for many secondary clock genes, including orthologs for vrille, slmb/lin23, shaggy/GSK3, and doubletime/Ck1e/KIN-20 (Additional file 15: Fig. S12). However, none of the orthologs or other homologues (Additional file 1: Table S11 & 12) have diel expression, even in the male head samples. Although *S. mansoni* orthologs of metazoan clock genes did not show 24-hour periodicity, we found 51 diel genes in *S.mansoni* whose ortholog in another animal also has ∼24 hour oscillations in expression (Additional file 1: Table S4 & 13). *S. mansoni* has 24 diel genes in common with *Drosophila melanogaster* (14 genes for male worms, *p* = 0.03927; 15 for females, *p* = 0.000687), 32 with mouse (23 for male *p* = 0.5179; 17 for females *p* = 0.6248), and only 4 with another lophotrochozoan, the limpet *Cellana rota* (Schnytzer *et al*., 2018) (4 for females *p* = 0.01117; 2 for males *p* = 0.3675).

## Discussion

Growing evidence is revealing the importance of biological rhythms in parasites as a strategy to optimize survival within a host and transmission between hosts (Reece *et al*., 2017). Behavioural patterns have been recorded for decades (Westwood *et al*., 2019 and references therein), but only in the last five years, in unicellular parasites, have the underlying molecular oscillations that are responsible for these rhythms been investigated (Rijo-Ferreira *et al*., 2017; Rijo-Ferreira *et al*., 2020). In this study, we have discovered the first 24-hour rhythms in the transcriptomes of a metazoan parasite; revealing that, like free-living animals, the effects of the earth’s daily rotation influences the biology of this intravascular flatworm, indirectly, at least. Our study also provides the first insights into biological rhythms of adult *S. mansoni*. We have demonstrated that the transcriptomes of this parasite are not static over the daily cycle, however, the number of diel genes is low compared to other animals; e.g. 5.7% in the water flea, *Daphnia pulex* (Rund *et al*. 2016) and 43% of protein coding genes in mouse show circadian rhythms, largely in an organ-specific manner (Zhang *et al*., 2014). We believe there are two reasons for this. First, we pooled six whole animals from a mouse, and sampled three mice at each time point, so any variation in rhythms between host mice, between worms within a pool (from one mouse), and between tissues within each worm, could mask organ-specific oscillations. We expect that many more diel genes will be identified as specific cell types and organs are sampled at this temporal scale, facilitated by recent advances in single cell transcriptomic methods (Wang *et al*., 2018; Wendt *et al*., 2020; Diaz *et al*., 2020).

Second, we carried out an initial filtering step to exclude genes from our analysis that were not significantly differentially expressed over a 24 hour period. The JTK_Cycle method alone does not determine whether a transcript varies significantly over time. So this filtering step has reduced false positives, and enabled us to identify diel genes that cycle with high enough amplitudes to be detected in pooled, whole worm samples. The amplitudes of diel genes were also low compared to other animals e.g. median fold change of 2 in *D. pulex* (Rund *et al*. 2016); as diel genes in male head samples had a higher median amplitude than the whole-body samples this suggests that organ-specific cycling might have been dampened in our whole animal samples. The effects of daily oscillations in transcripts has implications for all future RNAseq and functional genomic studies in this species, and the identification of diel genes in this study will allow future experiments to take into account, and control for, the effects of oscillating transcripts.

We interpreted the daily rhythms in the transcriptomes of *S.mansoni* within the context of what is known about the daily oscillations in mouse vasculature (Fig. 6). The putative function of *S. mansoni* diel genes and their cell type expression suggest that a number of host daily rhythms may be important zeitgebers to the worms’ rhythms; i.e. heat shock triggers, immune and coagulation factors and insulin. The most striking 24-hour rhythm was the nocturnal increase in expression of genes involved in the unfolded protein and stress responses. This revealed rhythmicity in molecular chaperones that are implicated in a wide variety of cellular processes; stabilizing new proteins (Tissières *et al*. 1974), processing proteins damaged by environmental stressors (Rampelt *et al*., 2012), and cell signalling (Stice & Knowlton 2008). Chaperones that have 24-hour periodicity in other animals (and are orthologs of *S. mansoni* diel chaperones) are controlled by the circadian clock to regulate protein aggregation and toxicity (Xu *et al*., 2019), and maintain the unfolded protein response (UPR) within a physiologically appropriate range (Eletto *et al*., 2014). In *S. mansoni*, increased UPR activity at night might be a response to, or anticipation of, periods of high ER-protein-folding demand that could be triggered by host cyclical rhythms, and this may be an adaptive stress response. One possible trigger is the daily body temperature cycle of the mouse, which increases by up to 4°C as it becomes active and starts feeding at the onset of the dark phase (Damiola *et al*., 2000). Although a relatively moderate increase, this is enough to activate the mouse’s own heat shock response (Reinke *et al*., 2008). The daily body temperature cycles of the mouse also drive a rhythmic alternative splicing (AS) program in itself (Preusner *et al*., 2017). It is a possibility, therefore, that the increase in the mouse body temperature at night also drives a heat shock response and splicing activity in the worms. Several RNA-binding proteins are involved in splicing regulation in response to heat stress; Ser/Arg-rich splicing factors are known regulators after heat shock (Shin *et al*., 2004), some hnrpH genes have a role in an arrest of mRNA splicing following heat shock (Honore, 2000; Mahe *et al*., 1997), and Hsp70 is known to reactivate mRNA splicing after heat inactivation (Vogel *et al*., 1995). Orthologs of these genes in *S.mansoni* form part of the night-time interaction network in male worms (Fig. 2). As temperature entrains circadian rhythms in *in vitro* populations of the blood-dwelling unicellular parasite, *Trypanosoma brucei* (Rijo-Ferriera *et al*., 2017), it may be an important cyclical cue for *S.mansoni* as well. Other environmental stressors known to activate the heat shock pathway include hypoxia and reactive oxygen species (Kregel, 2002) and as mouse blood oxygen and glucose levels increase during the dark phase (Adamovich *et al*., 2017; Llanos & de Vacarro, 1972), they could also induce the transcription of HSPs in *S. mansoni*. Circumstantial evidence points towards the nocturnal increased expression of heat shock and related genes being involved in proteotoxic stress and hormone cell signalling rather than a period of protein synthesis. This is because it is in anti-phase (at the opposite time of day) to rhythmic processes involved in translation regulation and protein synthesis, and it is synchronous with the mouse’s active phase and accompanying increases in environmental stressors.

The most prominent day-time process was a single extended metabolic ‘rush hour’ that started at the beginning of the hosts resting phase. As feeding can act to gate the initiation of metabolic activities (Sonoda *et al*., 2007) this may indicate a period of nutrient uptake in the worms. *S. mansoni* adults take up glucose and amino acids from the host blood directly across the tegument (Skelly *et al*., 2014), and some diel metabolic genes are markers of tegumental cell types. Worms also ingest blood cells into the digestive system with females thought to feed continuously, and males intermittently (Skelly *et al*., 2014), but there was no evidence of daily rhythms in genes involved in blood cell feeding. Insulin signalling, activated by host insulin, plays an important role in the growth, development and fecundity of schistosomes (Du *et al*., 2017), and is known to influence the maturation of schistosome eggs and their movement into the intestine (You *et al*., 2012; You *et al*., 2015). In mice, blood insulin levels peak at night (Feillet *et al*., 2016, Fig. 6). In the worms, throughout the night, transcript abundance of genes involved in the insulin signalling pathway increased, and so did egg-laying. It is possible, therefore, that the worms’ increase in egg laying rates coincide with periods of higher host insulin levels. However, because this egg-laying experiment was carried out *in vitro*, and therefore in the absence of host cyclical cues, this pattern must be an endogenous rhythm. Host cues, like insulin, however, could act as zeitgebers to synchronise the parasites rhythms to that of its host.

Although elements of the circadian clock network are conserved across diverse animal lineages (Hardin, 2011; Takahashi, 2017), the more taxa that are investigated the greater the variation discovered (Bell-Pedersen *et al*., 2005; Perrigault & Tran, 2017; Cook *et al*., 2018). A daily program of gene expression clearly exists in *S. mansoni* despite the lack of most of the canonical core clock gene orthologs. This suggests there is either an unusual oscillatory mechanism, or a functional endogenous clock has been lost and the *S. mansoni* rhythms are responding directly to host rhythms. The only core clock gene we found in the genome was Timeout, but it didn’t show 24-hour periodicity in expression. While its paralog, *Timeless,* functions as a canonical circadian clock gene in *Drosophila* and some other insects (Zheng & Sehgal, 2008; Iwai *et al*., 2006; Zhu *et al*., 2008), *Timeout* is a multifunctional gene. Indeed, *Timeout* plays an essential role in the maintenance of chromosome integrity, light entrainment of the circadian clock, embryonic development and regulation of DNA replication (Benna *et al*., 2010; Gotter *et al*., 2000; Gotter *et al*., 2007), as well as a role in the mammalian circadian clock (Barnes *et al*., 2003). It has circadian rhythms in the free-living flatworm *Schmidtea mediterranea* (Tsoumtsa *et al*., 2017); and, in female parasitic fig wasps, it only becomes rhythmically expressed once the wasp has successfully dispersed from the dark cavity of the fig, suggesting that rhythmicity is light-dependent (Gu *et al*., 2014). The last common ancestor of the Bilateria is hypothesized to have had all core clock components; Period, Timeless and Timeout, Clock, Cycle/Bmal1 and Cryptochromes (Reitzel *et al*., 2010), and combinations of these are present in extant Lophotrochozoa (Zantke *et al*., 2013; Perrigault & Tran, 2017; Cook *et al*., 2018). This suggests that *S. mansoni* has either lost all but *Timeout*, or these genes have diverged beyond recognition. *S. mansoni* orthologs of secondary clock genes did not show 24-hour periodicity. This could be a sampling artefact due to sequencing RNA from pooled, whole worm samples as clock gene rhythmicity may be limited to a subset of tissues (Whitmore *et al*., 1998), or the phase of clock gene expression can vary between tissues and even between cells within a tissue (Escamilla-Chimal *et al*., 2010; Wen *et al*., 2020). However, even in the male head samples (containing a subset of organs and tissues) there was still no 24-hour cycling of any putative clock gene transcripts. Alternative explanations for lack of 24-hour periodicity could be that they do not have rhythmicity at the transcript level, and/or may have non-clock functions. The adaptive advantage of a clock in environments with neither light nor very high-amplitude environmental cycles is less obvious (Olmedo *et al*., 2012). If, however, in future studies, any of these daily rhythms are discovered to be endogenous circadian rhythms, then our findings suggest that the *S. mansoni* clockwork must be quite distinct from that in other animals, and novel endogenous oscillators may be discovered within our list of diel genes.

Despite the profound global impact of schistosomiasis, there is complete reliance on only a single drug (praziquantel) for treatment, and evidence of reduced susceptibility in some schistosome populations (Ismail *et al*., 1996; Crellen *et al*., 2016), raises the spectre of drug resistance rendering current control measures ineffective. Consequently, there is a drive to develop a new generation of therapeutics based on schistosome genomes and their function (e.g. Berriman *et al*., 2009; Crosnier *et al*., 2020; Wang *et al*., 2020). Understanding the rhythms of target genes and their products will determine how an organ, or organism, will respond to a drug at a specific time of the day, and the timing of drug delivery could have a large impact on the effectiveness of target activation or inhibition (Cederroth *et al*., 2019). An RNAi screen to uncover new therapeutic targets in *S. mansoni* identified 195 genes that caused parasite detachment and affected survival (Wang *et al*., 2020), eight of which we have identified as diel genes (Additional file 1: Table S4), including *Hsp90* (Smp_072330) that demonstrated very high amplitudes in both sexes. By searching the ChEMBL database (Mendez *et al*., 2019), we identified existing drugs that are predicted to target the encoded protein of 26 diel genes, 12 of which are phase IV approved drugs (i.e. with the best safety record for humans), including four metabolic chokepoints (Additional file 1: Table S14). The diel genes with the highest amplitudes in each dataset are all putative drug targets, for example *SmKI-1* (Smp_307450) has four phase IV compounds that are predicted to target it, and it is also a proposed vaccine candidate (Hernandez-Goenaga *et al*. 2019). Therefore our fine temporal scale analyses of the *S. mansoni* transcriptomes will provide a useful foundation for the development and delivery of new therapeutics. Although we have described daily rhythms in schistosomes collected from nocturnal mice, we can assume that some of these rhythms will be inverted in worms infecting diurnal humans. The development of new therapeutics against schistosomiasis should include chronobiological information from the parasite and host wherever possible, and investigating further the temporal periods of parasite vulnerability (e.g. the stress response) and metabolic chokepoint activity (during the metabolic rush hour) holds promise for improving human health.

## Conclusions

This study has advanced our understanding of daily molecular oscillations in organisms to now include a metazoan parasite. Schistosome adults live in the bloodstream of a mammalian host, which is a 24-hour rhythmic environment. Our finding that *S. mansoni* adults have daily rhythms in their transcriptomes is, therefore, not entirely unexpected. These daily rhythms in the parasite may be driven by host rhythms, either directly, and/or generated by an intrinsic circadian clock that is entrained to host cues. What is surprising, however, is that exploration of the genome revealed a lack of core clock genes that are generally conserved across other animals, and this is suggestive of an unusual oscillatory mechanism or loss of a functional endogenous clock. Our findings provide a crucial first step from which future studies can examine how these rhythmic accumulations of mRNA abundance; i) are generated; i.e. the balance between synthesis and degradation; ii) are propagated, i.e. imposed by the host versus endogenously controlled by *S.mansoni* itself; and iii) if they eventually exert functions. Most importantly, our identification of diel genes and daily processes has revealed fine-scale temporal partitioning of biological processes, some of which may serve the particular time-of-day challenges of life within the host; e.g. the activity of the hosts immune and haemostatic systems. These rhythmic genes and processes give us insight into how these parasites can survive for decades in this environment, and highlight the need to incorporate daily oscillations in transcript abundance into functional genomic studies aimed at developing and delivering novel therapeutics against schistosomiasis.

## Methods

### Animal procedures

The life cycle of *Schistosoma mansoni* NMRI (Puerto Rican) strain is maintained at the Wellcome Sanger Institute (WSI) by breeding and infecting susceptible *Biomphalaria glabrata* snails and mice. Female Balb/c mice were bred at the WSI, and maintained on individual air handling units at 19 to 23°C and 45–65% humidity. Animals were given access to food and water ad libitum, maintained on a 12-hour light/dark cycle, and housed in groups of no more than 5 adults per cage. Welfare assessments are carried out daily, abnormal signs of behaviour or clinical signs of concern are reported. All personnel at the WSI performing welfare checks on animals are trained and assessed as competent by qualified named individuals.

Thirty-six 6 weeks old female were percutaneously infected with 200 mixed-sex *Schistosoma mansoni* cercariae collected from 13 infected snails as described (Crosnier *et al*., 2019). In brief, under isoflurane anaesthesia, the mice were carefully transferred onto individual holders in a bespoke pre-warmed anaesthesia rig and their tails inserted into the test tubes containing with the cercariae. After 40 minutes exposure, animals are removed from the anaesthesia rigs, placed back into their cage and monitored until full recovery from the anaesthesia. For parasite collection (below) mice were euthanised by intraperitoneal injection of 200 µl of 200 mg/ml pentobarbital (Dolethal®) supplemented with 100 U/ml heparin (cat.# H3393, Sigma Aldrich), and adult worms recovered by portal perfusion (the portal vein is sectioned followed by intracardiac perfusion with phenol-red-free DMEM, cat.# 31053-044 ThermoFisher Scientific, containing 10 U/mL heparin)

### Parasite collection

At 42 days post infection, groups of 3 mice were perfused every 4 hours for 44 hours, and the adult worms collected. The worms sampled in the dark phase were collected from mice euthanized under red light conditions. We collected worm samples 30 minutes after lights on (Zeitgeber time (ZT) 0.5) and then 4 hours subsequently giving us collection times of ZT:0.5, 4.5, 8.5, 12.5, 16.5, 20.5 over two 24hr periods. ZT:0.5 corresponds to 8am in the human 24hr clock, so actual collection times were 08:00, 12:00, 16:00, 20:00, 00:00, 04:00. At each collection time worms were washed in serum-free DMEM media at 37°C. Mature, paired male and female worms from each mouse were separated and six female worms were pooled and stored in TRIzol at -80°C, and the same for six male worms. A further 10 male worms were pooled from each mouse and fixed and stored in the RNA stabiliser *vivo*PHIX^TM^ (RNAssist Ltd, Cambridge, UK) at 4°C for the dissection of heads the following week. We used male heads only as they are bigger and easier to dissect than female heads. RNA was extracted from each pool and sequenced (**Figure 1A**). From each mouse at each time point, we therefore collected material simultaneously for 3 time-series datasets: pooled females, pooled males and pooled male heads, with 36 samples in male and female datasets, and 33 samples of male heads (three perfusions had too few worms to collect for head samples: day1_20:00_b, day2_04:00_b, day2_04:00_c).

### RNA isolation, library preparation and transcriptome sequencing

RNA was isolated from the pooled whole worm samples in TRIzol reagent according to the manufacturer’s instructions (Life Technologies). For the male head samples, ten additional male worms per mouse per time point were dissected in *vivo*PHIX^TM^ by cutting posterior to the ventral sucker and anterior to the testes. The heads were rinsed in 50% ethanol and pooled in TRIzol and the RNA extracted as for the whole worm samples. RNA quality was assessed using a Pico RNA kit for the BioAnalyzer (Agilent). We were able to extract good quality RNA from all but one sample (male head day2_20:00_c). Total RNA was enriched for mRNA using poly(A) pulldown. The sequencing libraries were prepared using the NEB Ultra II RNA custom kit on an Agilent Bravo WS automation system. All samples had 14 cycles of PCR, which was set-up using Kapa HiFi Hot start mix and Eurofins dual indexed tag barcodes on Agilent Bravo WS automation system. RNA sequencing of the pooled worm libraries was performed on six lanes of the Illumina HiSeq2500 v4 75 Paired End sequencing platform. All sequencing data are available through ENA study accession number ERP108923.

### Identification of diel cycling transcripts

Read quality was assessed using the FASTQC quality control tool. Raw reads were mapped to the *Schistosoma mansoni* genome (version 7, WormBaseParaSite (WBPS version 14, WS271)) using STAR (Dobin, 2013). Genes with fewer than 10 reads mapping across all samples were excluded. Read counts were normalised using DESeq2 v1.22.2 (Love *et al*., 2014) with default parameters and the variance stabilising transformation. Principal Components Analysis was then used to exclude outlier samples by comparing them to replicates. One male sample (day1_12:00_a), two male head samples (day1_20:00_b; day2_20:00_a) and no female samples were excluded.

To reduce false positive calls of cycling genes, we first excluded genes which were not differentially expressed across the time course using the GLM approach in edgeR v3.24.3 (Robinson *et al.,* 2010). Initially only genes with Counts Per Million (CPM) greater than three across at least three samples were included in the analysis. In the model design replicates for equivalent Circadian time in each of the two 24h periods were considered as the same time point. Genes with a False Discovery Rate (FDR) greater than 0.05 were then excluded.

To identify cycling genes, we used JTK_cycle (Hughes *et al.,* 2010), called using the meta2d function from the MetaCycle software (Wu *et al.,* 2016). Default parameters were used i.e. minimum period 20 hours, maximum period 28 hours. Genes with an FDR < 0.01(JTK BH.Q <0.01) were called as cycling. For visualisation of cycling gene expression, normalised counts for replicates were averaged and then log-transformed to generate heatmaps using the *pheatmap* package with the option scale = ‘row’. The fold change of gene expression over the time points was calculated as the max (peak)/ min (trough), keeping replicates separate.

### Identification of drug targets

Drugs from the ChEMBL database that we predict to interact with the cycling genes were identified using the approach described in Wang *et al*. (2020).

### Gene Ontology enrichment analysis of cycling genes

To better understand the function of genes identified as cycling, we performed Gene Ontology (GO) enrichment analysis of our gene lists using topGO (Alexa *et al*., 2006), with FDR < 0.05, node_size = 5, method = ‘weight01’, statistic = ‘Fisher’. GO terms for *S. mansoni* were downloaded from WormBase Parasite using BioMart (Howe *et al*., 2016) on the 5th March 2020.

### KEGG pathway mapping

Mapping of *S. mansoni* gene products to the KEGG pathway database was performed on the KAAS server (https://www.genome.jp/kegg/kaas/) using the GHOSTX program and BBH method. The significance of cycling gene enrichment in pathways was assessed using Fisher’s Exact test and resulting P-values were adjusted using the Benjamini-Hochberg procedure, where maps in the KEGG categories 1-4 (https://www.genome.jp/kegg/pathway.html) were tested. Pathways with FDR < 0.05 were considered as significant. For visualisation the R package Pathview (Luo *et al*., 2013) was used.

### Molecular interactions analysis

Molecular interactions were predicted using the online search tool STRING (www.string-db.org; V 11)(Szklarczyk *et al.,* 2015). The *S. mansoni* V7 gene identifiers for diel gene were converted to *S. mansoni* V5 gene identifiers. The protein sequences for V5 gene identifiers were analysed in STRINGdb. Protein sequences for day-time peaking genes for each dataset were entered as a multiple protein search. Default settings were used to predict interactions with a minimum interaction (confidence) score of 0.4, corresponding to medium level of confidence. A second identical analysis was carried out for night time peaking genes in male and female worms.

### Single cell data analysis

We used publicly available single cell transcriptome data from mature adult male and female worms (Wendt *et al*., 2020) to determine if cycling transcripts were specific to a certain cell type (i.e. cell type markers), specific to a category of cell types (e.g. muscle, neurons, germline, tegument lineage etc), or more broadly expressed and found in more than one category of cell type. Processed single-cell RNA-seq data were provided by the authors (R object Whole_Integrate_rmv27_50_RN.rds). We used the hypergeometric test to determine whether each cell type cluster contained more marker genes called cycling in our datasets than expected by chance. This was done separately for each of the male, female and male head datasets. The resulting p-values were corrected using the Benjamini-Hochberg method. A Python script implementing this method is available from our GitHub page (https://github.com/adamjamesreid/schistosoma_daily_rhythms/).

Seurat UMAP plots were used to explore the expression of cycling transcripts across cell types and whether they were ubiquitously expressed or enriched, or specific, to one or more cell type (Stuart *et al*., 2019). Some of the cycling transcripts identified as cell type markers and enriched in specific cell types were validated by *in situ* hybridization (below).

### Identification of hypothetical proteins

Amino acid sequences for the ten diel hypothetical proteins were obtained from WormBaseParasite. Protein 3D structures were predicted from amino acid sequences using I-TASSER online server(v5.0) (Yang *et al.,* 2015) with default parameters. TM-scores indicate similarity between two structures. The values range from 0-1, with the value of 1 indicating a perfect match.

### *Fluorescent in situ* hybridisation and imaging

Mature adult pairs, collected from mice infected for life cycle maintenance and parasite material production, were anaesthetised in 0.5% solution of ethyl 3-aminobenzoate methanesulfonate (Sigma-Aldrich, St. Louis, MO) for 15 minutes to separate male and female worms. The worms were killed in 0.6 M MgCl2 for 1 min, and incubated in 4% formaldehyde in PBSTx (1xPBS + 0.3% TritonX) for 4 hours at room temperature. They were rinsed 3 x 5minutes in PBSTx, dehydrated into 100% Methanol and stored at -20C. Samples were gradually rehydrated in PBSTx over 30minutes and incubated in 5ug/ml Proteinase K (Invitrogen) in 1x PBSTx for 30 minutes at 37°C. They were post-fixed in 4% Formaldehyde in PBSTx for 10 min at room temperature then rinsed in PBSTx for 10 minutes. Probes, buffers, and hairpins for third generation *in situ* hybridization chain reaction (HCR) experiments were purchased from Molecular Instruments (Los Angeles, California, USA). Experiments were performed following the protocol described by Choi *et al*. (2016; 2018) and developed for wholemount nematode larvae. Samples were mounted using DAPI fluoromount-G (Southern Biotech) and imaged on a confocal laser microscope (Sp8 Leica).

### *In vitro* egg laying assay

Twelve freshly perfused pairs of adult worms (still coupled) were placed into individual wells of a 12-well plate containing 3 ml of ABC169 media (Wang *et al*., 2019) and kept at 37°C, 5%CO2 in the dark. Eggs from each well were collected, and counted, at 8am and 8pm every day for 72 hours, giving 3 day-time counts and 3 night-time counts per worm couple. The first 12 hour period post-perfusion was discounted to allow the worms to acclimate to the *in vitro* conditions. This experiment was replicated 3 times, each time with freshly perfused worms. The median egg number for each worm for day-time, and night-time was calculated and a paired Wilcoxon test was carried out to determine if there was a significant difference in the number of eggs laid between day or night.

### Identification of core, and secondary, circadian clock genes in *Schistosoma mansoni*

We identified putative *Schistosoma mansoni* homologues of animal circadian clock genes using two methodologies. The first was a BLASTP sequence similarity search with a cut off e-value of 1e-10 against the *S. mansoni* genome (v7) in WormBaseParaSite (WBPS version 14, WS271), using previously defined circadian proteins sequences from UniProtKB (Boutet *et al*.2007) and GenBank (**Supplementary Table 9**). Our second method enhanced the robustness of our searches by using respective domains of proteins to identify putative orthologues, as shown before (Padalino *et al*. 2018). Briefly, domain identifiers for main clock proteins were selected using Pfam and SMART, and their respective signatures were used to query the BioMart function in WBPS against the entire *S. mansoni* genome (**Supplementary Table 10**). A BLASTP of output sequences in NCBI (Altschul *et al*.1990) was used to identify these proteins, and all respective hits were aligned in Jalview 2 and illustrated in IBS illustrator (Liu *et al*. 2015).

To examine whether hits were orthologous to circadian clock proteins from other animals, phylogenetic analyses on the core clock components were conducted; Timeless/Timeout, Cycle/BMAL1/Arntl and Clock (all are basic helix-loop-helix-PAS proteins) and the Cryptochromes/ Photolyases, and the secondary clock proteins; Vrille, Slmb, Shaggy and Doubletime. Sequences were aligned using CLUSTAL OMEGA (Sievers *et al*. 2011) and visually examined using Jalview 2 (Waterhouse *et al*. 2009). The aligned sequences were exported into Gblocks 0.91b (Castresana, 2000) with allowance for smaller blocks and less strict flanking positions for reduced stringency. Conserved positions (3% for bHLH/PAS, 9% for CDP photolyase, 16% for timeless) were used to construct a Neighbour-Joining phylogenetic tree (JTT model) with partial/pairwise deletion and 1000 bootstrap replications in MEGA-X (Kumar *et al*. 2018).

### Comparison of diel 1-to-1 orthologs in *Schistosoma mansoni* and other Metazoa

We identified 2925 cycling genes in *Drosophila melanogaster* from the Cycling Gene Data Base (CGDB; Li *et al*., 2017) and 3233 *S. mansoni*-*D. melanogaster* one-to-one orthologues from Wormbase Parasite (Bolt *et al*., 2018). Of the one-to-one orthologues, 420 cycled in *D. melanogaster*, 66 in *S. mansoni* males, 48 in females. For mouse, we identified 9534 cycling genes from CGDB and 2855 one-to-one orthologues with *S. mansoni* using Wormbase Parasite. 1146 shared orthologues were cycling in mouse, 57 in male, 44 in female. To examine common cycling genes between *S. mansoni* and another lophotrochozoan, we used the 221 cycling limpet (*Cellana rota*) transcripts identified by Schnytzer *et al*. (2018). A total of 38,482 limpet translated transcript sequences were used with 14499 sequences from *S. mansoni* (WBPS15) to identify one-to-one orthologues using OrthoFinder (Emms & Kelly, 2019). Here we looked for shared orthogroups rather than one-to-one orthologues due to the fragmented nature of the limpet transcriptome assembly. There were 5025 shared orthogroups between limpet and *S. mansoni*. 67 limpet cyclers and 96 *S. mansoni* cyclers were in shared orthogroups. We used the Fisher exact test to determine whether the number of one-to-one orthologues cycling in both species was greater than expected by chance.

## Declarations

### Ethics approval and consent to participate

The procedures involving animals were conducted under the Home Office Project Licence No. P77E8A062 held by GR. All protocols were revised and approved by the Animal Welfare and Ethical Review Body (AWERB) of the WSI. The AWERB is constituted as required by the UK Animals (Scientific Procedures) Act 1986 Amendment Regulations 2012.

### Consent for publication

Not applicable

### Availability of data and materials

The datasets generated and analysed during the current study are available in the European Nucleotide Archive (www.ebi.ac.uk/ena) under the ENA study accession number ERP108923.

### Authors’ contributions

KAR, PD, GR, CM, AJR and MB conceived and designed the sequencing experiments. KAR, PD, GR, CM, GS collected the material and prepared it for sequencing. NEH and MS coordinated the sequencing. AJR, ZL, PD, AC and KAR analysed the sequencing data. KAR, PD, ZL, AJR, CD, SKB, GR and MB interpreted the sequencing data. AW, KH and DW identified and analysed circadian clock gene orthologs. The paper was prepared by KAR and all authors read and approved the final manuscript.

## Acknowledgements

We thank all members of the Parasite Genomics team at the Wellcome Sanger Institute for their comments and input on this study. We thank the animal facility staff – Simon Clare and his team, and David Goulding of the Electron and Advanced Light Microscope Facility for their support & assistance in this work. We thank Andrew Goldsborough of RNAssist Ltd for advice on RNA stabilisation and Anna Protasio for discussion on RNA sequencing. We thank Jim Collins and his lab for helpful discussion on their single cell data. The infrastructure used for this analysis is maintained by the core IT Service and the Pathogen Informatics teams at Wellcome Sanger Institute.

## Supplementary figures

**Supplementary Fig. S1.**
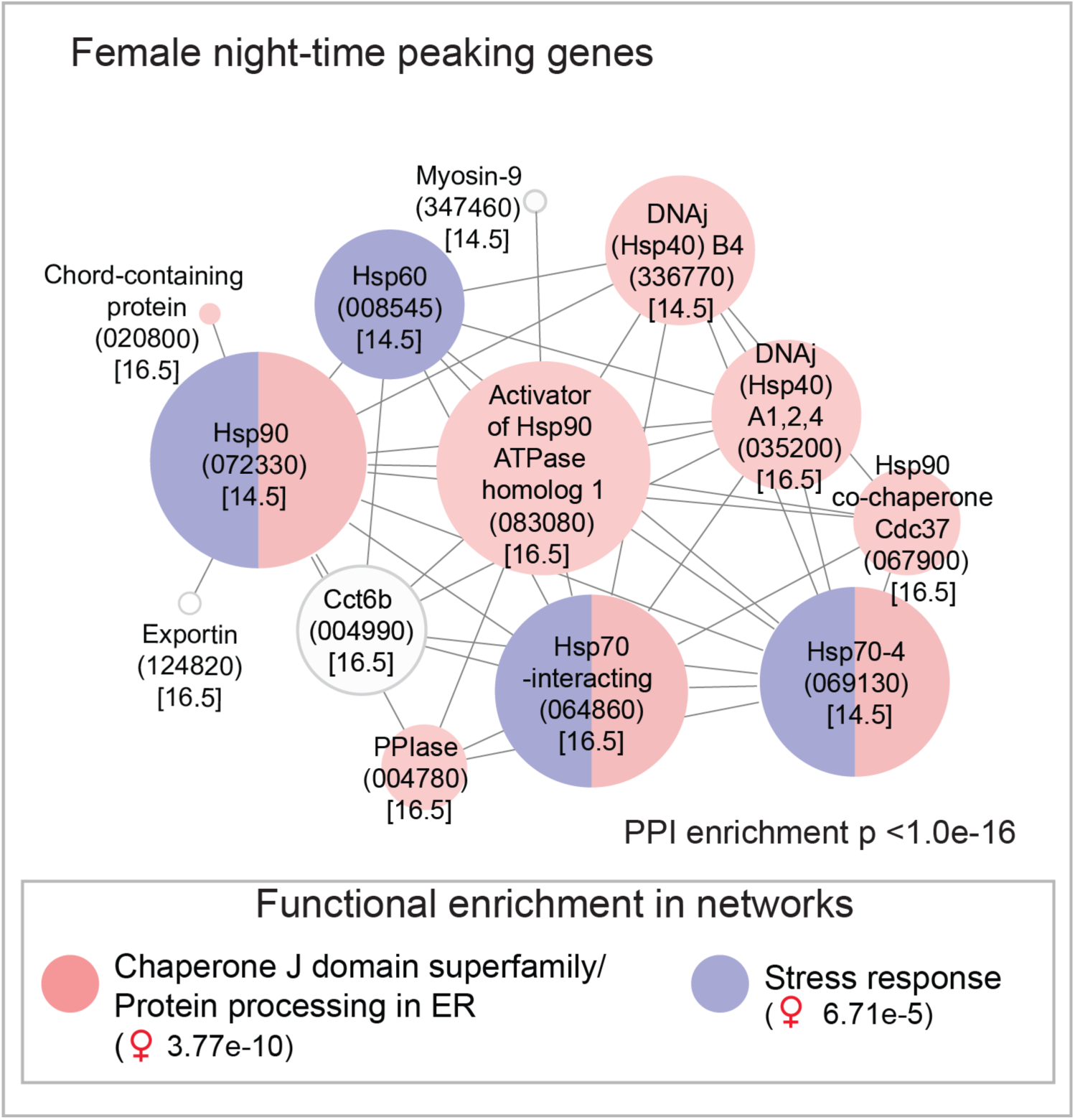
Predicted molecular interaction networks of night-time peaking genes in female *Schistosoma mansoni* (computed using the STRING online database). Node size reflects the number of connections a molecule has within the network. Lines (edges) connecting nodes are based on evidence of the function of homologues. Functional enrichment (FDR) as provided by STRING. (PPI= predicted protein interaction; geneIDs with Smp_ prefixes removed; acrophase in brackets).

**Supplementary Fig. S2.**
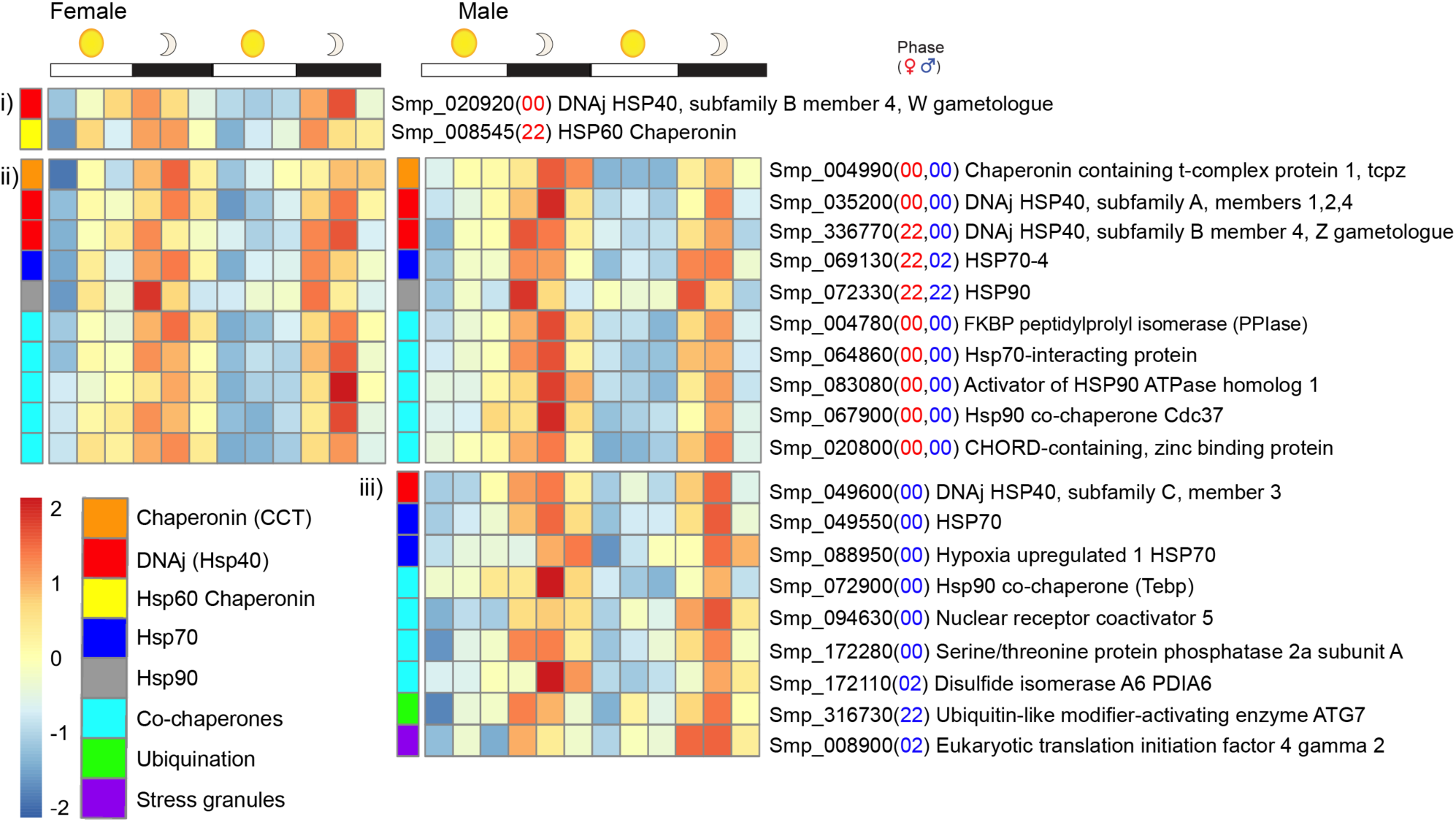
Diel genes encoding heat shock proteins, co-chaperones and other proteins involved in heat shock response and recovery. All reach their acrophase between 22:00-02:00 hours but some show diel expression in one sex only (i & iii), whereas another ten cycle in both sexes, with eight in phase (ii).

**Supplementary Fig. S3.**
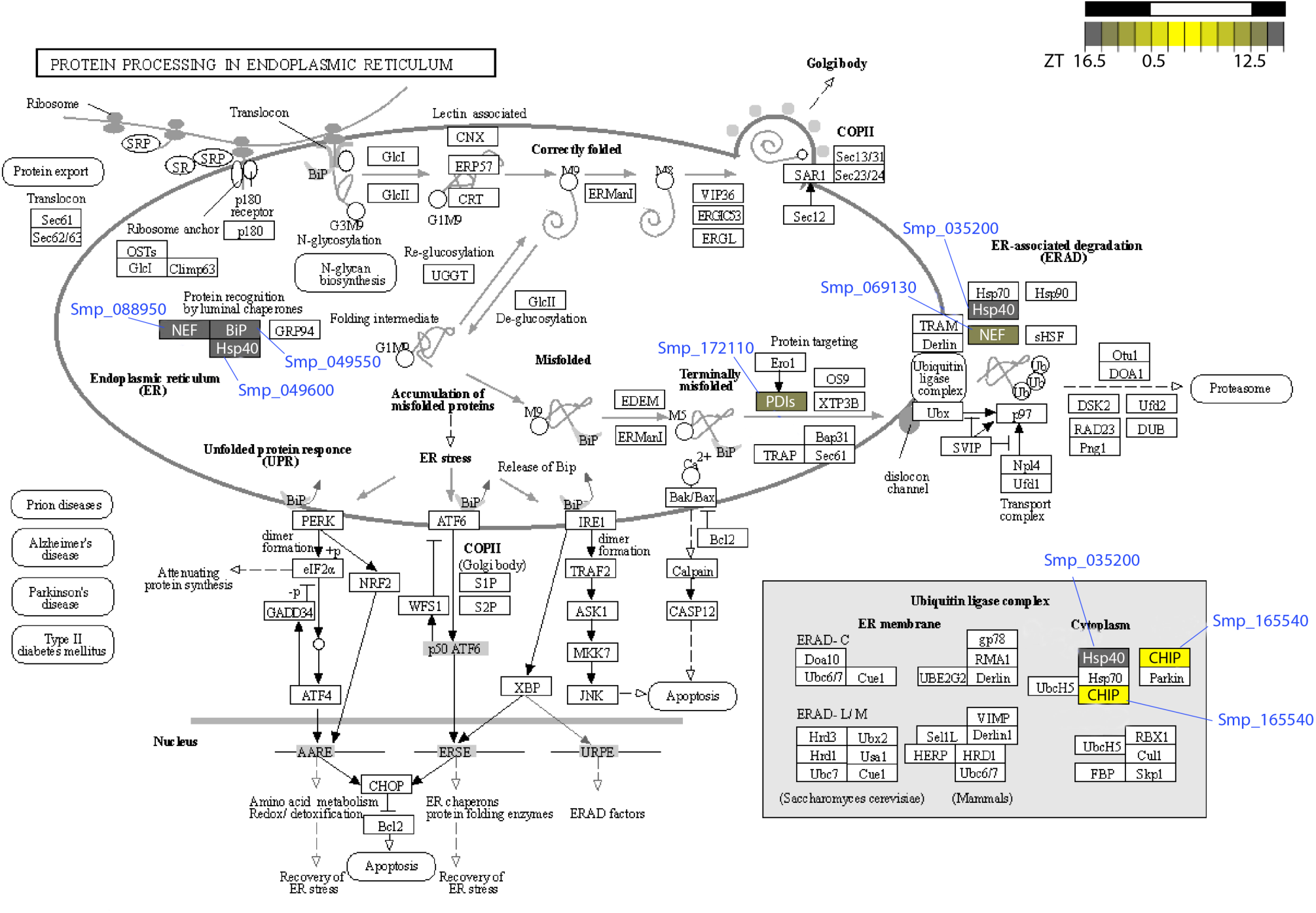
The KEGG pathway ‘Protein processing in endoplasmic reticulum’ includes seven diel genes that encode heat shock proteins and other co-chaperones. Data on KEGG graph rendered by Pathview.

**Supplementary Fig. S4.**
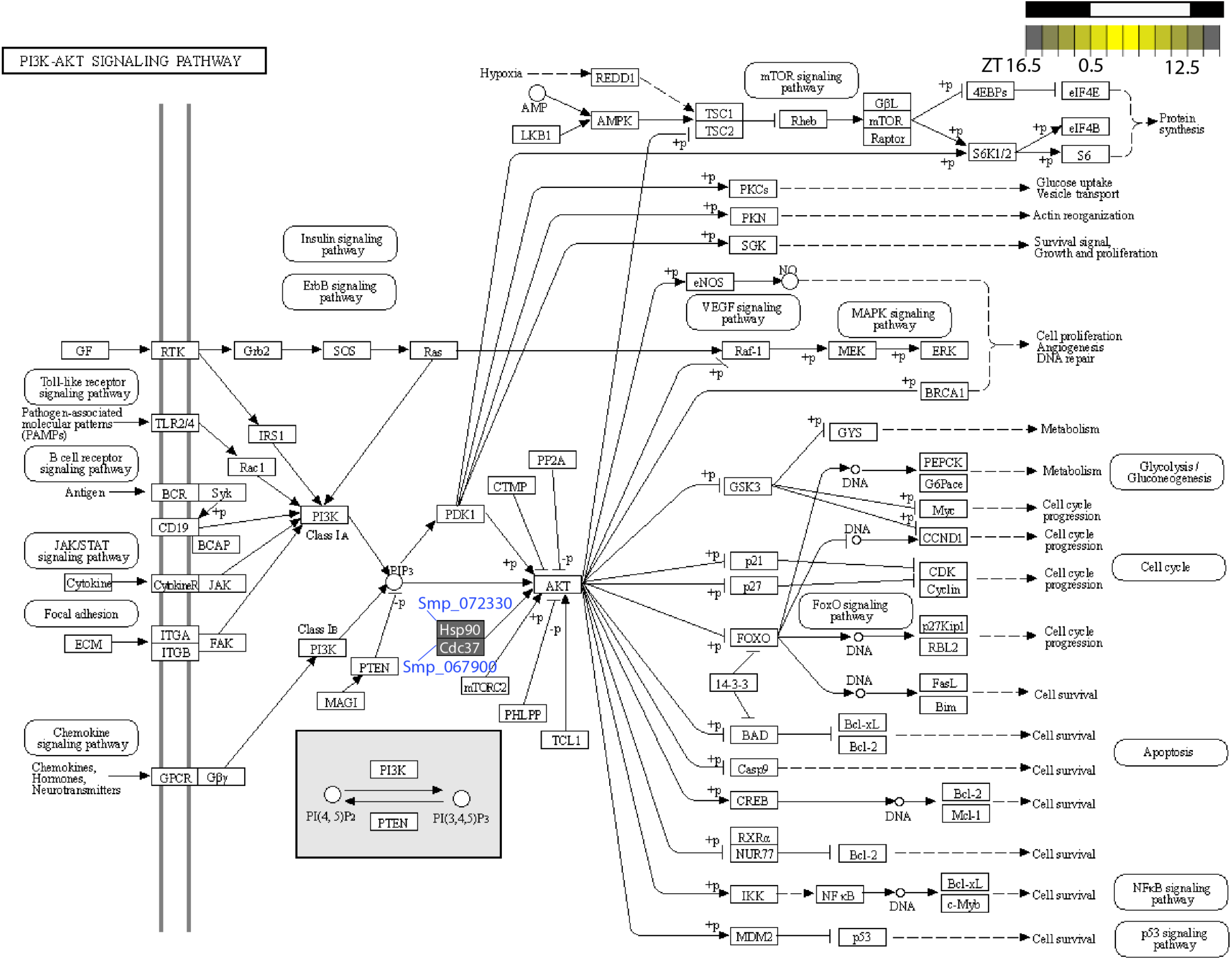
The KEGG pathway ‘PI3K-AKT signaling pathway’ includes two diel genes; one that encodes heat shock protein 90 (HSP90) and the other encodes one of its co-chaperones, cell division cycle 37 (Cdc37). Data on KEGG graph rendered by Pathview.

**Supplementary Fig. S5.**
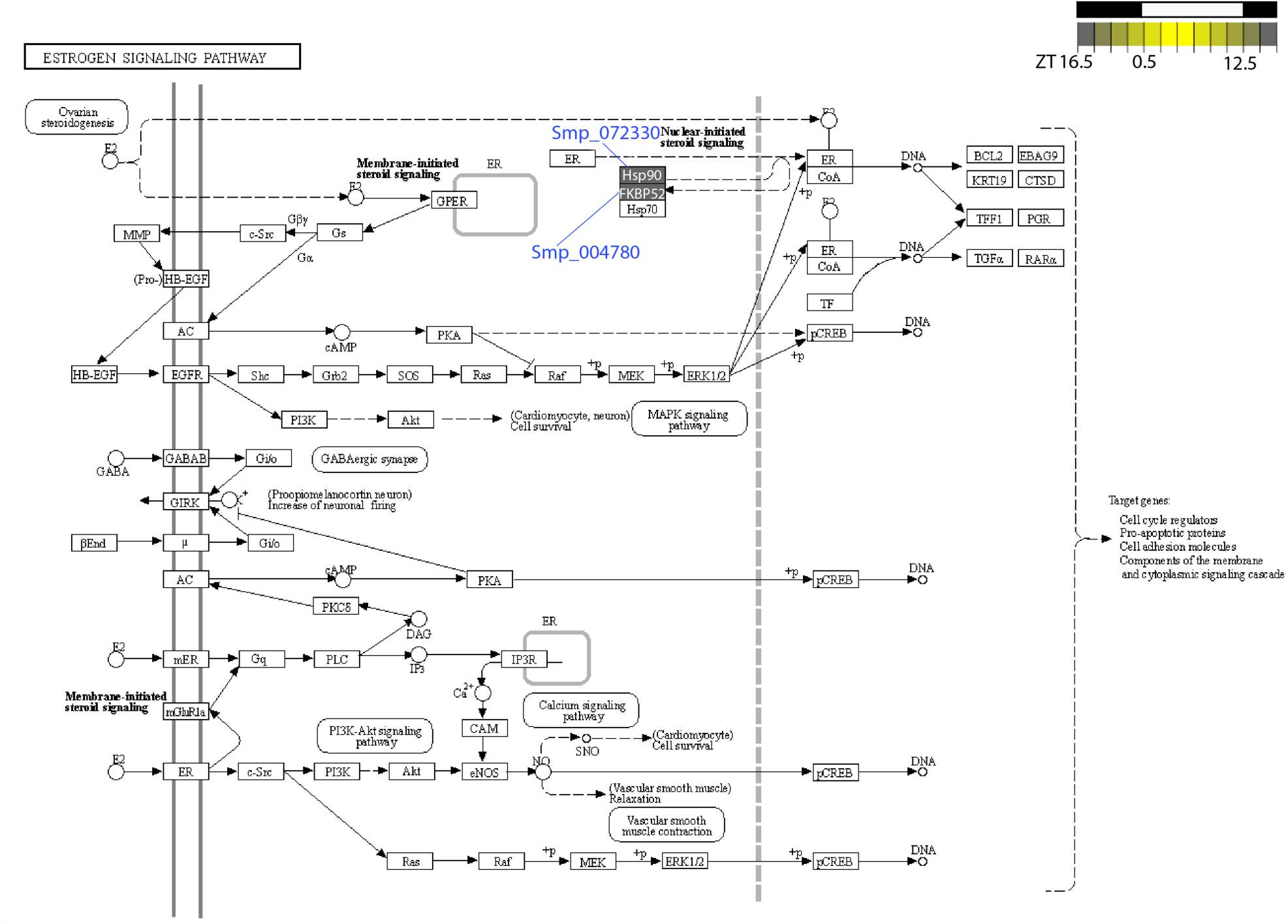
The KEGG pathway ‘Estrogen signaling pathway’ includes two diel genes; one that encodes heat shock protein 90 (HSP90) and the other encodes one of its co-chaperones, immunophilin (FKBP52). However, HSP70 (Smp_303420), another binding partner, does not cycle in male or female worms. Data on KEGG graph rendered by Pathview.

**Supplementary Fig. S6.**
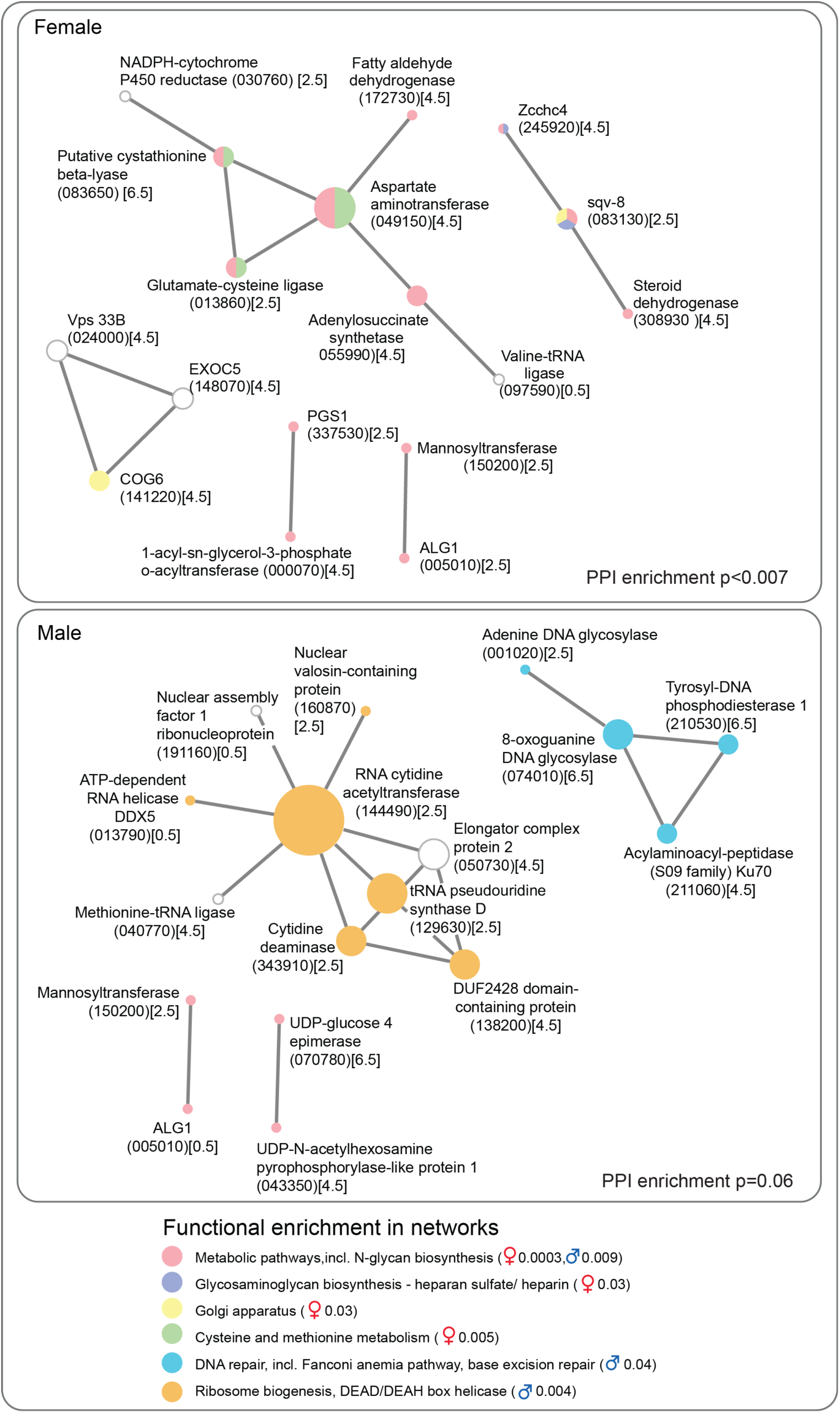
Predicted molecular interaction networks of day-time peaking genes in female and male *Schistosoma mansoni* (computed using the STRING online database). Node size reflects the number of connections a molecule has within the network. Lines (edges) connecting nodes are based on evidence of the function of homologues. Functional enrichment (FDR) as provided by STRING. (PPI= predicted protein interaction; “Smp_” prefixes have been removed from gene identifiers for clarity; acrophase in brackets).

**Supplementary Fig. S7.**
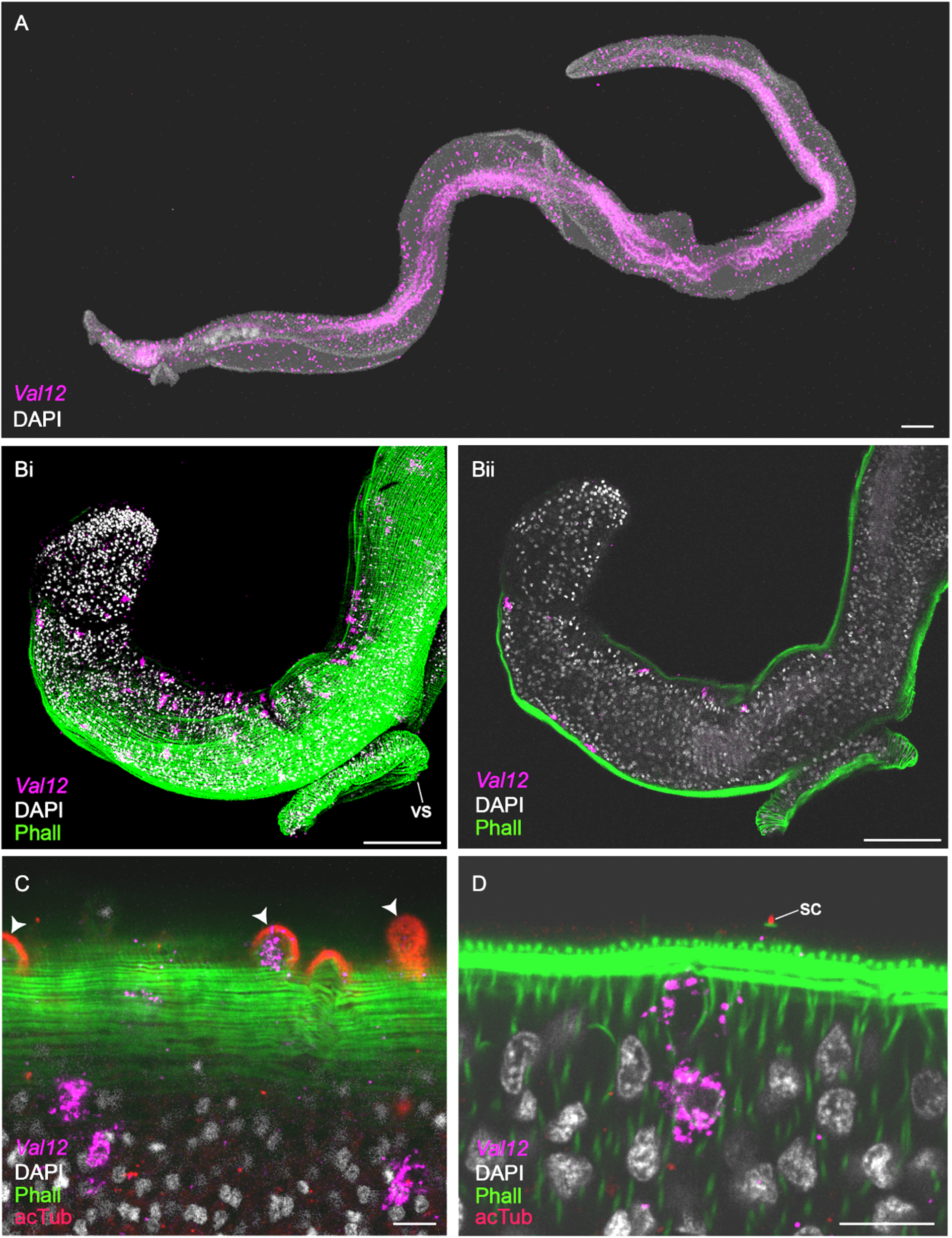
In male worms *Venom allergen-like 12* (*Val12*, Smp_123540) transcripts peak in abundance at midday (ZT 4.5) and are expressed in **A)** ∼ 500 cells that are distributed the length of the body (anterior to left)(scale = 200µm). **B)** The *Val12*^+^ cells sit below the body wall musculature (phall = phalloidin) on both the ventral and dorsal sides, **i**) 3D projection and **ii**) optical section of the head (scale = 100µm). **C&D**) *Val12* is expressed in some tubercles (arrowhead) of the dorsal tegument, as well as directly under the body wall musculature (scale = 10µm)(optical sections)(acTub = acetylated tubulin).vs = ventral sucker, sc = sensory cilia. 100% of individuals examined, n = 20.

**Supplementary Fig. S8.**
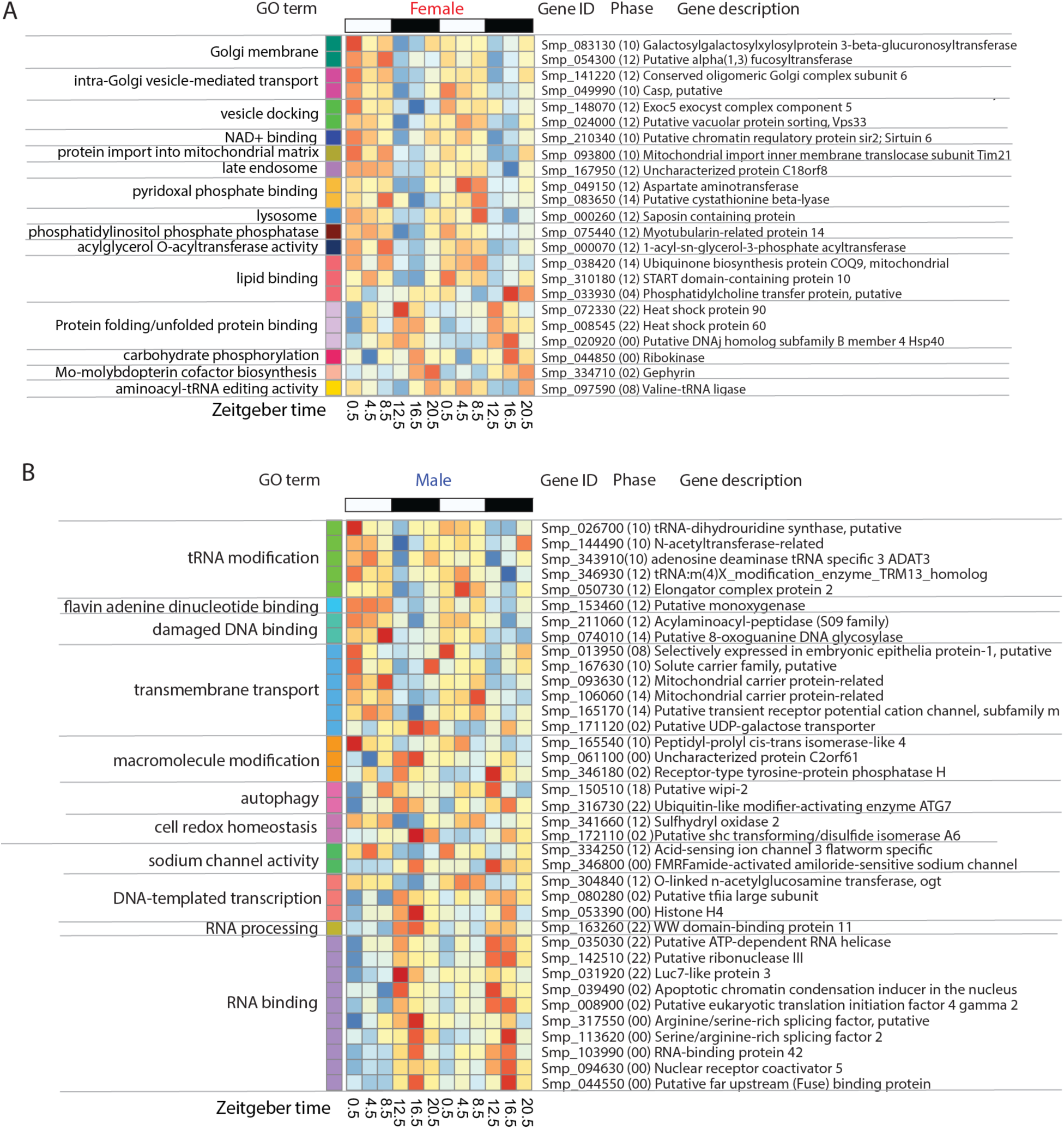
Sex-specific 24-hour rhythmic processes. Heatmaps showing GO terms enriched in diel genes that cycle in females (**A**) or males (**B**) only.

**Supplementary Fig. S9.**
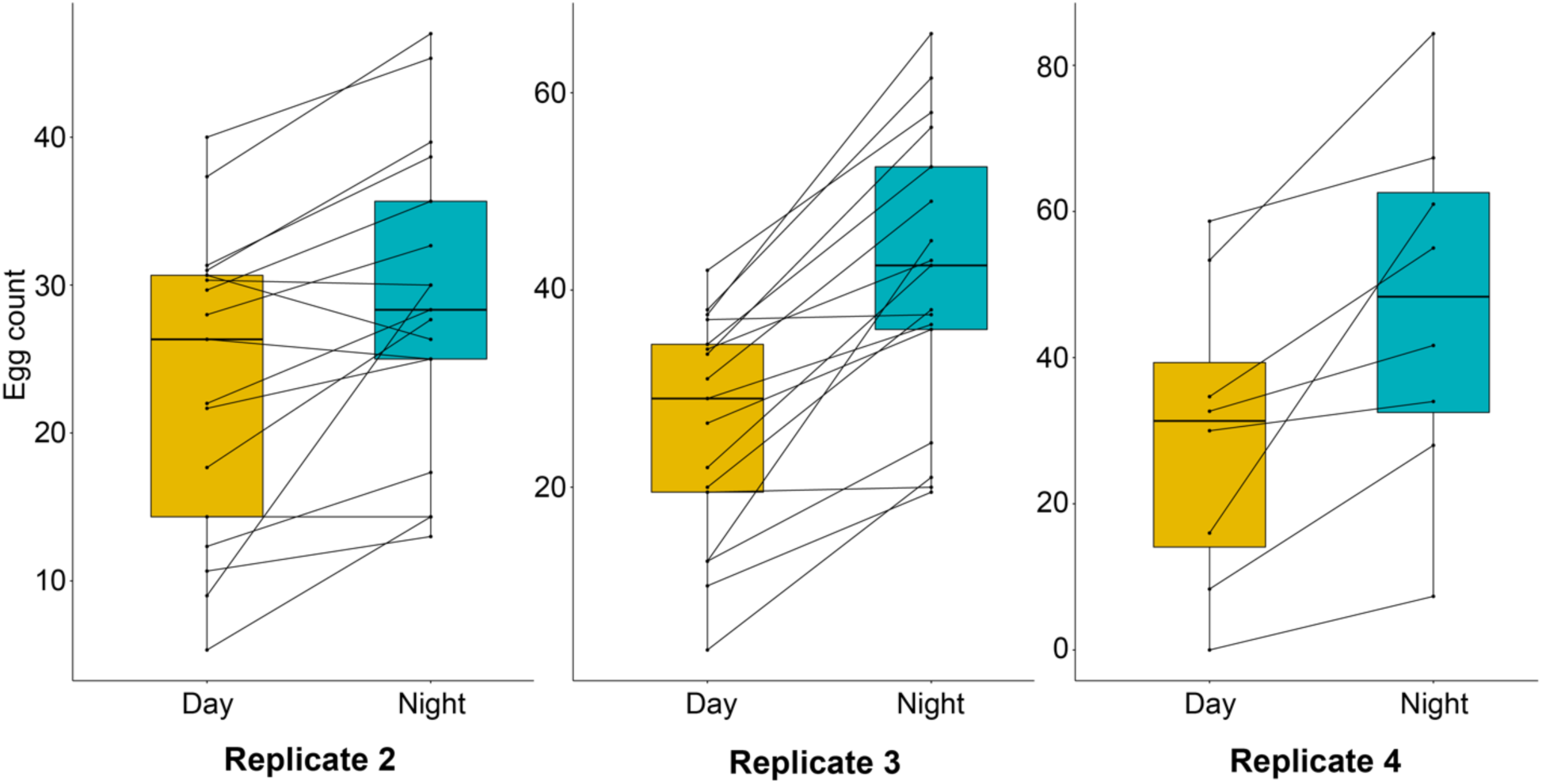
Independent biological replicates of day and night egg counts from paired female worms *in vitro* (median and interquartile ranges). Replicate 2: n=17, paired Wilcoxon test P=0.002, median(night-day)=5.3. Replicate 3: n=17, P=0.0003, median(night-day)=17.5. Replicate 4: n=8, P=0.008, median(night-day)=14.3.

**Supplementary Fig. S10.**
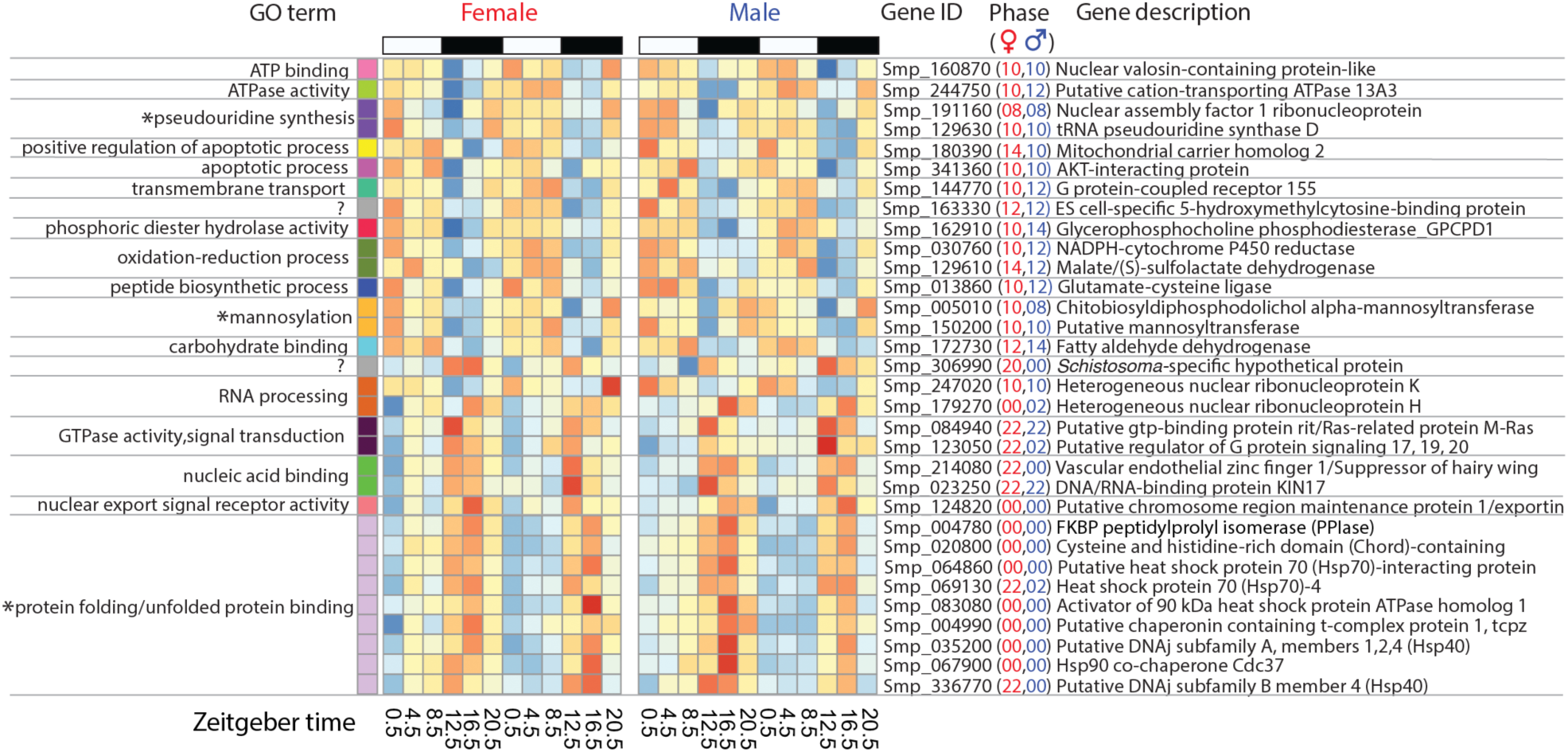
Diel genes common to female and male worms show identical, or similar, phases suggesting that many biological processes and molecular functions (GO terms) are happening in synchrony. Most enriched functions are time-of-day specific; e.g. mannosylation, redox homeostasis and apoptosis occur during the daytime, whereas genes involved in molecular chaperoning, nucleic acid binding and signal transduction reach their acrophase at night. (* significantly enriched GO terms FDR<0.01, **supplementary table 5**).

**Supplementary Fig. S11.**
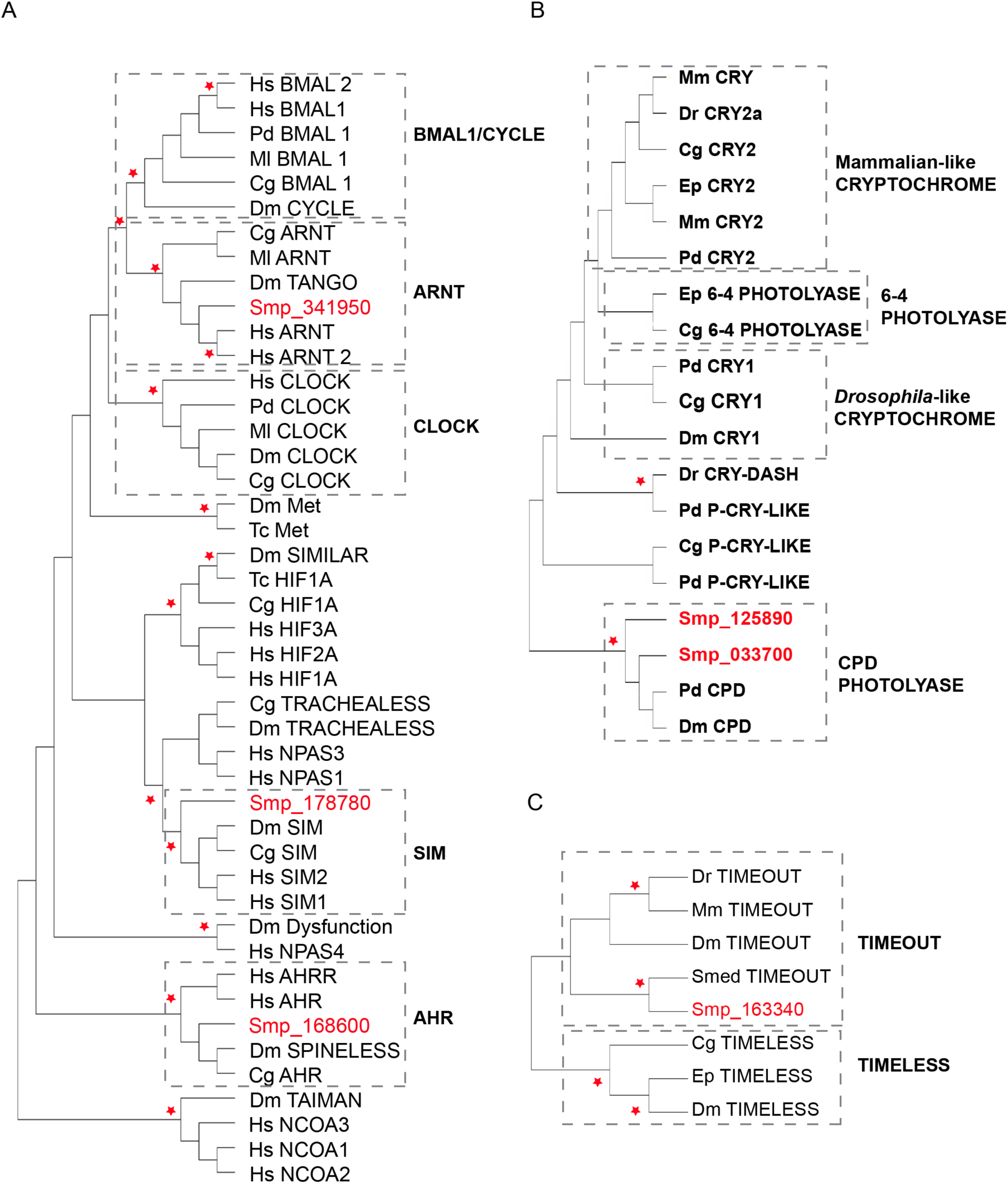
Core circadian clock gene homologs are missing in *Schistosoma mansoni*. Neighbour-joining phylogenetic trees were constructed in Mega-X with 1000 bootstrap replications and partial/pairwise deletion (* = bootstrap support >90). **A**) A phylogeny of bHLH/PAS domain proteins shows that *S.mansoni* lacks BMAL1 and CLOCK homologs. Our BLASTP hits cluster with the closely-related non-circadian proteins ARNT, AHR and SIM. **B**) A phylogeny of orthologous genes *tim1* (TIMELESS) and *tim2* (TIMEOUT) shows that our BLASTP hit clusters with *tim2*. We also show that the previously identified *timeless* homolog in the flatworm *Schmidtea mediterreana* (Tsoumtsa et al., 2017) clusters with *tim2* homologs of model organisms. **B)** *S. mansoni* has two CPD photolyases but no circadian-related Cryptochromes. Abbreviations: Cg = *Crassostrea gigas*, Dr *= Danio rerio,* Dm = *Drosophila melanogaster,* Ep = *Eurydice pulchra*, Hs = *Homo sapiens,* Ml = *Melibe leonina*, *Mm = Mus musculus,* Pd = *Platynereis dumerilii*, Smed = *Schmidtea mediterranea*, Tc = *Tribolium castaneum*

**Supplementary Fig. S12.**
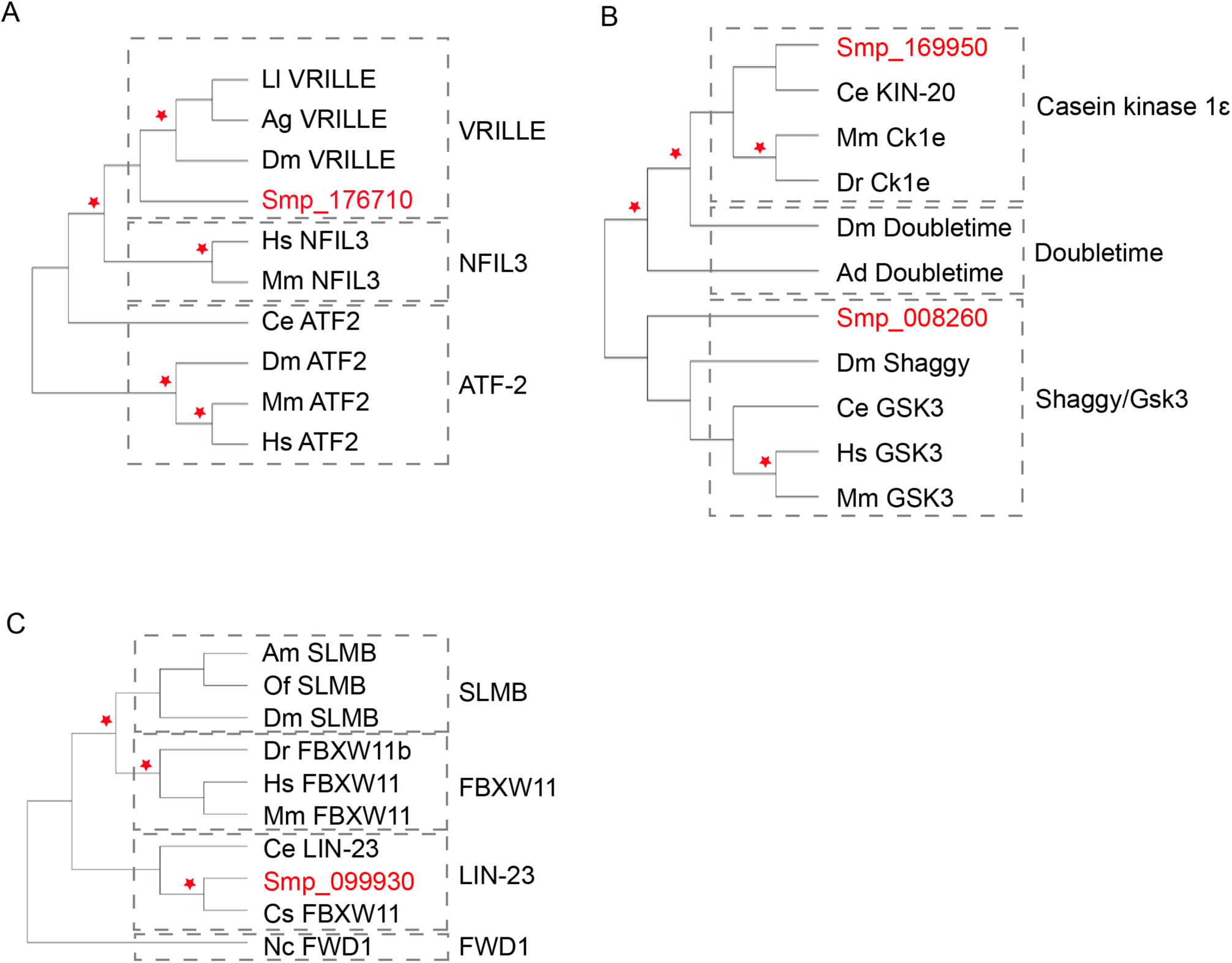
*Schistosoma mansoni* has homologs of secondary clock genes. Neighbour-joining phylogenetic trees were constructed in Mega-X with 1000 bootstrap replications and 80% partial deletion using sequences obtained from Uniprot. (* = bootstrap support >90). **A**) Representatives of basic-leucine zipper protein family show insect *vrille* homolog is present in *S. mansoni*. **B**) Smp_169950 clusters with mammalian homologs of *doubletime*, whereas Smp_008260 clusters with *Shaggy*. **C**) Smp_099930 clusters with Lin-23, a previously identified homolog of Slmb in *Caenorhabditis elegans*. Ancestral sequence from *Neurospora Crassa* was used as an out- group. Conserved regions were obtained in Gblocks using least stringency criteria and percentage of all sequences used was as follows: 3% basic-leucine zipper, 11% slmb, 18% shaggy/doubletime. Abbreviations: Ll = *Lutzomyia longipalpis*, Ag = *Anopheles gambiae*, Dm = *Drosophila melanogaster*, Hs = *Homo sapiens*, Mm = *Mus musculus*, Ce = *Caenorhabditis elegans*, Cg = *Crassostrea gigas,* Am = *Apis mellifera*, Of = *Oncopeltus fasciatus*, Dr *= Danio rerio*, Nc = *Neurospora crassa*, Cs = *Clonorchis sinensis*, Ad = *Anopheles darlingi*.

