## Additional file 8: supplementary info 1 for "Daily rhythms in the transcriptomes of the human parasite *Schistosoma mansoni*"

**Additional file 8: Supplementary information 1**

**Smp_307450 - SmKI-1, a Bovine Pancreatic Trypsin Inhibitor/Kunitz protease inhibitor domain protein**

Diel gene Smp_307450 was previously identified as Smp_147730 in version 5 of the genome. Smp_147730 has been split into 4 genes in version 7; Smp_307450, Smp_311660, Smp_311670, Smp_337730. Aligning Smp_147730 (from UniProt) to Smp_307450, Smp_311660, Smp_311670, Smp_337730, shows that they are all very similar; containing the Kunitz domain, with its six conserved cysteine residues (Laskowski & Kato, 1980), the Kunitz family signature (Ranashige et al., 2015), and the same amino acid residue at the reactive P_1_ site (Krowarsch et al., 1999). The P_1_ site is the major determinant of the specificity of protease recognition by Kunitz inhibitors; typical trypsin inhibitors contain Arg (R) or Lys (K) (Krowarsch et al., 1999). Smp_307450 differs from Smp_147730 at seven sites, but the conserved Arg at the P_1_ site indicates that Smp_307450 is potentially a trypsin inhibitor.

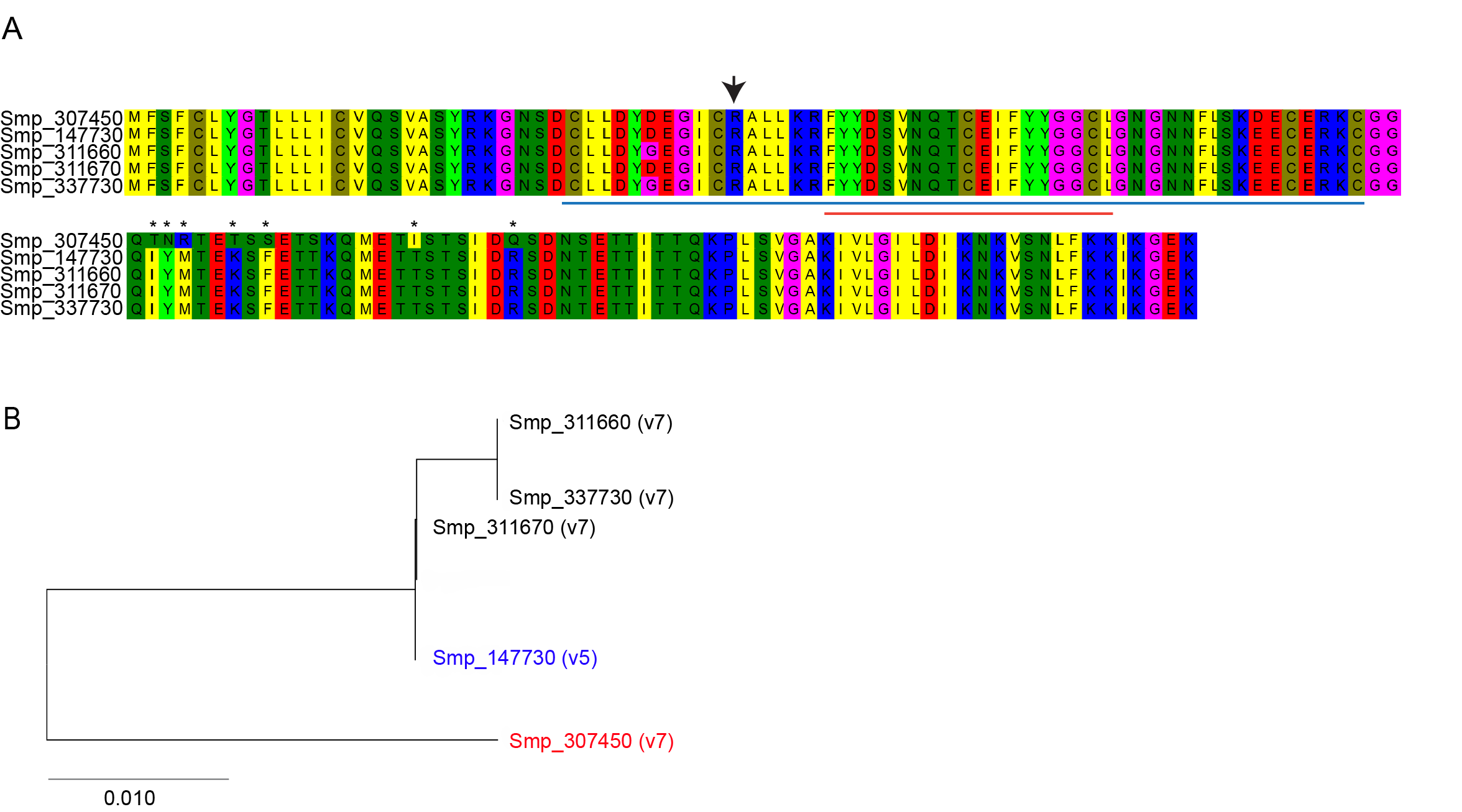

**Supplementary Information Figure 1. A)** ClustalW alignment and **B)** Neighbour joining tree (MEGA 7) of Smp_307450, Smp_311660, Smp_311670, Smp_337730 (v7 of genome) and Smp_147730 (v5). Kunitz domain (blue line), reactive P_1_ site (black arrow head), and the Kunitz family signature (red line). * amino acid differences between Smp_307450 and Smp_147730.
