## Additional file 14: supplementary info 2 for "Daily rhythms in the transcriptomes of the human parasite *Schistosoma mansoni*"

**Additional file 14: Supplementary information 2**

**Clock gene protein structure**

***Core clock genes***

**Basic-helix-loop-helix-PAS transcription factor family**

The basic-helix-loop-helix-PAS transcription factor family contains the domains bHLH, PAS, PAC (Wu & Rastinegad, 2017). CLOCK and CYCLE/BMAL1 are members that are known to be involved in the animal circadian clock (Fribourgh & Partch, 2017). CLOCK, in addition to the family-wide domains, has a characteristic poly-Q (polyglutamine) domain (SI Fig.2). Shortening of the poly-Q domain results in impairment of transcription activity of clock (Allada et al., 1998; Gesto et al., 2015). This domain is missing in our BLASTP hits (Smp_168600, Smp_178780 and Smp_341950) (SI Fig.2), which instead clustered within the AHR, SIM, and ARNT clades respectively (Supplementary figure 10). The identified ARNT homolog in *S. mansoni* (Smp_341950) also lacked the PAC motif in all three of its splice variants (SI Fig.2). The PAC motif contributes to PAS binding (Ponting & Aravind, 1997) and has been concluded to be a part of the PAS domain (Hefti et al., 2004).

**Timeless (tim1)/ Timeout (tim2)**

*tim1* and *tim2* are paralogous genes (Benna et al., 2000), with *tim1* duplicating from *tim2* at the time of the Cambrian explosion (Rubin et al., 2006). Like mouse and fly *tim2,* but unlike fly *tim1,* *S. mansoni,* and the free-living flatworm *S. mediterranea,* have *tim2* orthologues that contain a PAB domain at their C- terminus (SI Fig.2). In humans, the PAB domain is known to be involved in DNA repair by binding to a DNA repair enzyme Poly ADP-ribose polymerase 1 (PARP-1) (Xie et al., 2015).

**Cryptochrome/Photolyase family**

The FAD binding domain prominent in DNA Photolyases and Cryptochromes is absent in *S. mansoni’s* CPD photolyases (SI Fig.2).

***Secondary clock genes***

We identified an *S. mansoni* homolog of *Vrille* *(vri)* (Smp_176710). Whereas the insect model only has one Basic Leucine Zipper (bzip) domain, the *S. mansoni* homolog appeared to have a double bzip domain (SI Fig.2). The Activating Transcription Factor 2 (ATF-2) in *C. elegans* is the only current double bzip domain protein studied in literature (Cowell 2002; Wang et al., 2006) (Supplementary figure 11). Our hit for *doubletime* (Smp_169950) clustered with its mammalian homolog glycogen synthase kinase GSK3 (Supplementary figure 11). Neither SMART nor Pfam could identify an FBOX in our *slmb* homolog (SI Fig.2). However, our hit (Smp_099930) was clustered with *lin-23*, the *C. elegans* homolog of *slmb* (Supplementary figure 11).

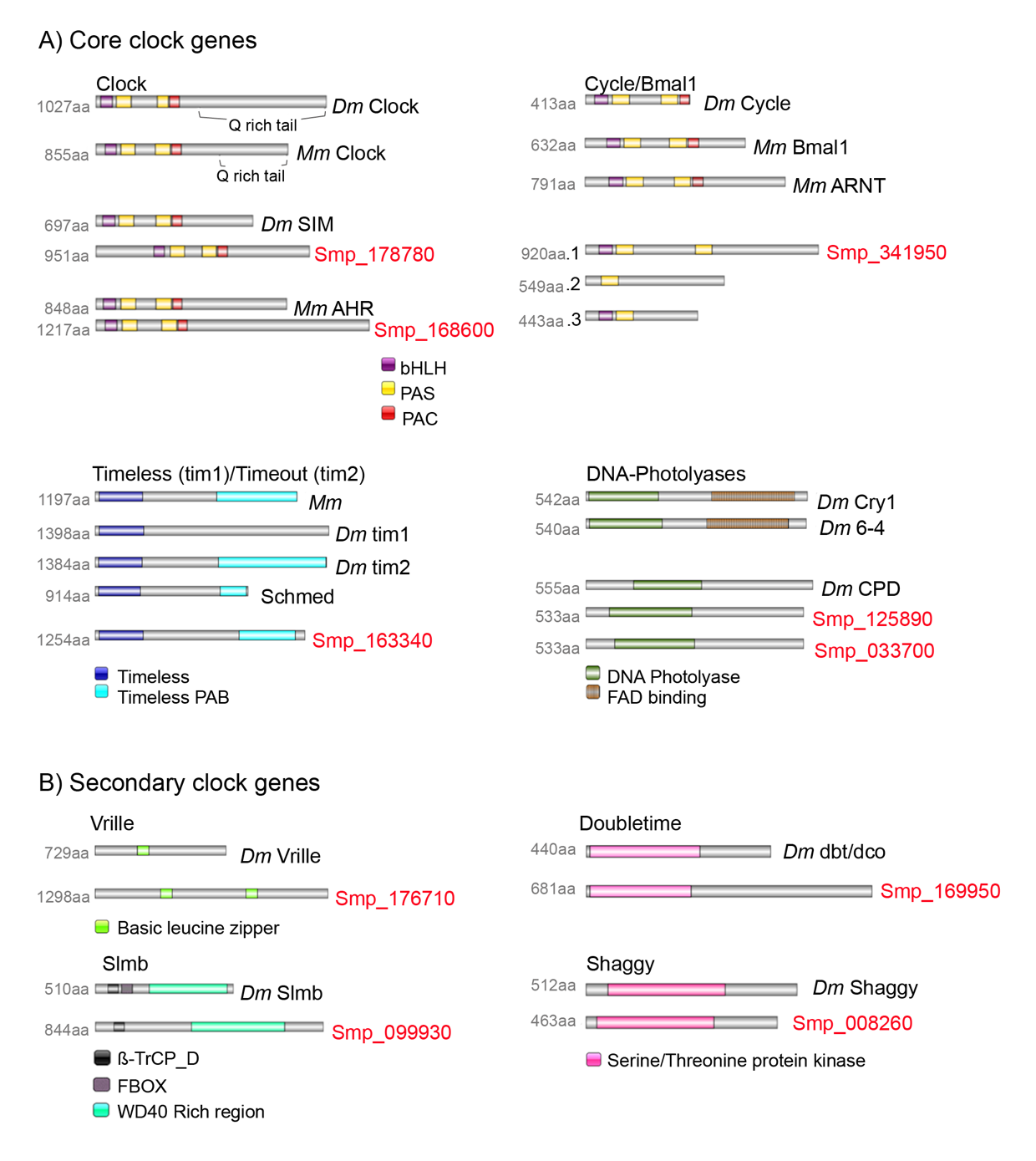

**Supplementary Information Figure 2. Secondary structural features of putative circadian proteins in *Schistosoma mansoni*.** **A)** Core clock components and **B)** secondary clock components. Note the missing Q-rich tail in SIM/ AHR and the missing FBOX domain in *Slmb* homolog. Amino acids are indicated on the left and names are shown on the right.
